# Shelterin and subtelomeric DNA sequences control nucleosome maintenance and genome stability

**DOI:** 10.1101/405712

**Authors:** Thomas van Emden, Marta Forn, Ignasi Forné, Zsuzsa Sarkadi, Matías Capella, Lucía Martín Caballero, Sabine Fischer-Burkart, Cornelia Brönner, Marco Simonetta, David Toczyski, Mario Halic, Axel Imhof, Sigurd Braun

**Affiliations:** Department of Physiological Chemistry, BioMedical Center (BMC), Ludwig Maximilians University of Munich, Martinsried, Germany; Protein Analysis Unit (ZfP), BioMedical Center (BMC), Ludwig Maximilians University of Munich, Martinsried, Germany; Department of Biochemistry, Gene Center, Ludwig Maximilians University of Munich, Germany; Department of Biophysics and Biochemistry, University of California San Francisco (UCSF), San Francisco, USA; International Max Planck Research School for Molecular and Cellular Life Sciences, Martinsried, Germany; Present address: Netherlands Cancer Institute, Department of Molecular Oncology, Amsterdam, The Netherlands

**Author notes:** These authors contribute equally to this work (Thomas van Emden, Marta Forn).

**Keywords:** telomeres, shelterin, heterochromatin, CLRC, SHREC, nucleosomes, genome stability

## Abstract

Telomeres and the shelterin complex cap and protect the ends of chromosomes. Telomeres are flanked by the subtelomeric sequences that have also been implicated in telomere regulation, although their role is not well defined. Here we show that, in *Schizosaccharomyces pombe*, the telo-mere-associated sequences (TAS) present on most subtelomeres are hyper-recombinogenic, have metastable nucleosomes, and unusual low levels of H3K9 methylation. Ccq1, a subunit of shelter-in, protects TAS from nucleosome loss by recruiting the heterochromatic repressor complexes CLRC and SHREC, thereby linking nucleosome stability to gene silencing. Nucleosome instability at TAS is independent of telomeric repeats and can be transmitted to an intrachromosomal locus containing an ectopic TAS fragment, indicating that this is an intrinsic property of the underlying DNA sequence. When telomerase recruitment is compromised in cells lacking Ccq1, DNA sequences present in the TAS promote recombination between chromosomal ends, independent of nucleosome abundance, implying an active function of these sequences in telomere maintenance. We propose that Ccq1 and fragile subtelomeres co-evolved to regulate telomere plasticity by controlling nucleosome occupancy and genome stability.

## INTRODUCTION

The chromatin of eukaryotic cells is organized into structural and functional domains that are crucial for genome stability. For example, centromeres and telomeres are composed of large repetitive DNA elements assembled into heterochromatin, which mediate the faithful inheritance and integrity of chromosomes. Telomeres are composed of G/T-rich tandem DNA repeats that provide docking sites for telomere-binding proteins to protect the chromosomal ends from erosion and fusion. In humans and *Schizosaccharomyces pombe,* telomere protection is mediated by the shelterin complex, which consists of six subunits (Fig. 1A): Pot1 (hPOT1); Tpz1 (hTPP1); Poz1 (hTIN2); Rap1; Taz1 (hTRF1 and hTRF2) (Cooper *et al*, 1997; Miyoshi *et al*, 2008; Ye *et al*, 2004; Nandakumar & Cech, 2013). Fission yeast has an additional subunit Ccq1, which associates with shelterin via Tpz1 and is essential for maintaining telomere length through recruiting telomerase (Armstrong *et al*, 2014; Flory *et al*, 2004; Harland *et al*, 2014; Miyoshi *et al*, 2008; Tomita & Cooper, 2008; Moser *et al*, 2015; Jun *et al*, 2013). When telomeres shorten to a critical size (after about 100 generations), cells undergo telomere crisis resulting in cell cycle arrest and cellular senescence (Dehé & Cooper, 2010). However, survivors can arise through telomerase-independent maintenance mechanisms. Cells without telomerase mainly survive through the formation of circular intrachromosomal fusions, whereas mutants lacking Ccq1 maintain linear chromosomes through a homologous recombination (HR) pathway (Tomita & Cooper, 2008).

**Figure 1.**
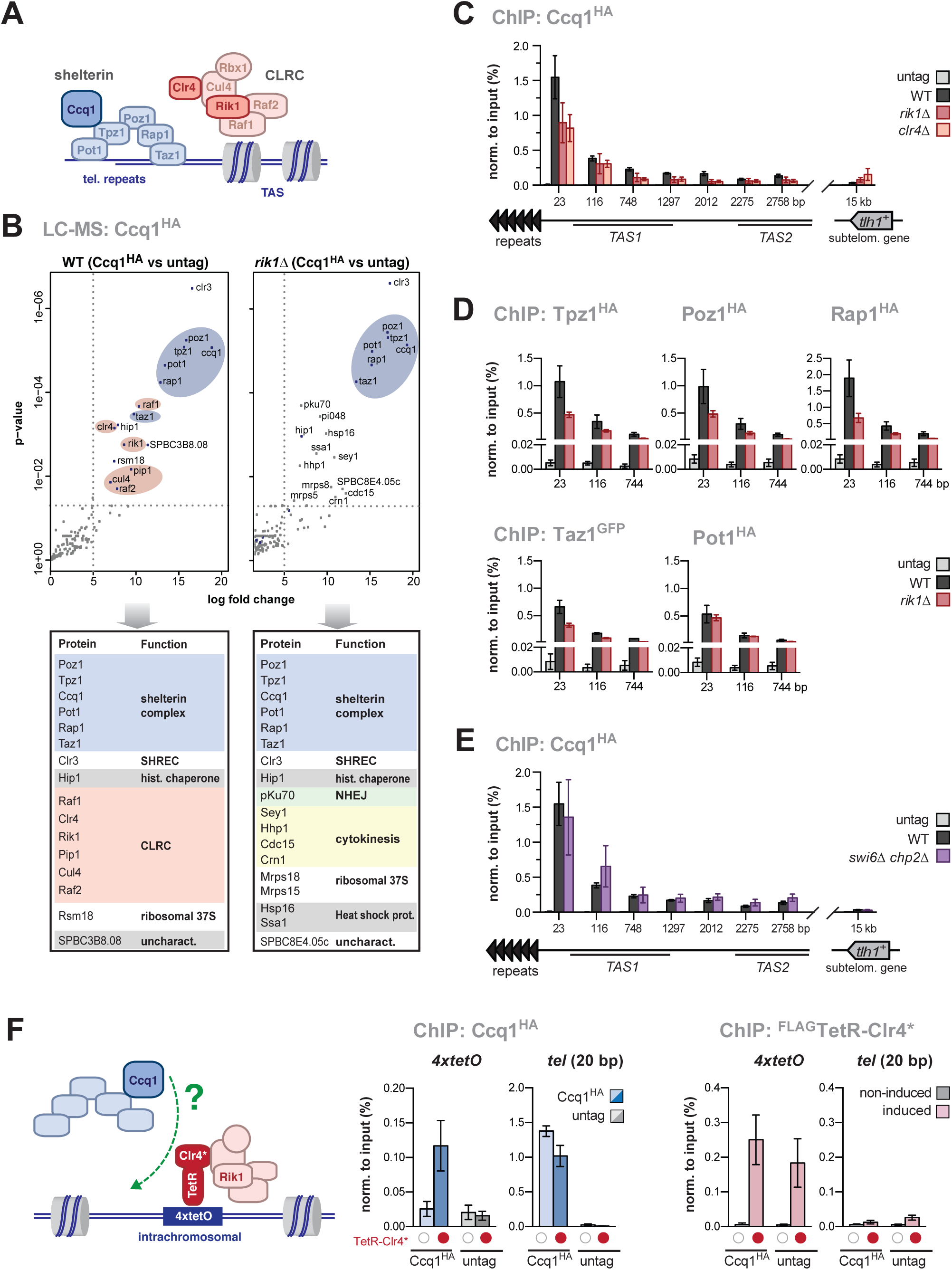
CLRC promotes the shelterin-chromatin association. **(A)** Scheme of the shelterin complex and CLRC at telomeres. **(B)** Mass spectrometry analysis of proteins co-purified with Ccq1^HA^. Shown is a volcano plot for proteins significantly enriched in Ccq1^HA^ relative to untagged Ccq1 (left panel: WT cells; right panel: *rik1Δ* cells). Members of the CLRC complex and shelterin are highlighted in red and blue, respectively. Bottom panel displays proteins enriched (log_2_ ≥ 5 or pval ≤ 0.01). **(C-E)** ChIP-qPCR analysis of epitope-tagged shelterin components in WT and mutant strains as indicated (negative control: untagged strain). Positions on x-axis denote distances relative to telomeric repeats (telomere-proximal PCR primer). **(F)** ChIP analysis of Ccq1^HA^ and FLAG-TetR-Clr4* (see scheme). Expression and tethering of FLAG-TetR-Clr4* are controlled by thiamine/AHT addition (red dots indicate induced/tethered FLAG-TetR-Clr4*; empty circles indicate non-induced/non-bound controls). For all ChIP analyses, data are normalized to input and represented as mean ± SEM from 3 independent experiments.

Subtelomeres often comprise homologous sequence elements that differ from the telomeric repeats, yet their function is not well defined. The subtelomeric homology (SH) sequences in *S. pombe* contain loosely repetitive elements known as telomere-associated sequences (TAS), which are enriched in poly[A/T] tracts and found telomere-proximal on nearly every chromosomal arm. Whereas SH sequences are dispensable for viability and telomere length, their absence causes harmful inter-chromosomal fusions in cells lacking telomerase (Tashiro *et al*, 2017), suggesting that they promote chromosome circularization upon telomere attrition. In addition, the subtelomeres may be involved in telomerase-independent, HR-mediated maintenance of linear chromosomes (Cooper *et al*, 1997; Nakamura *et al*, 1998); however whether these mechanisms require specific SH sequences remains unknown.

*S. pombe* subtelomeres harbor heterochromatic domains telomere-distal from the TAS, which are marked by high levels of di- and trimethylated lysine-9 of histone H3 (H3K9me2/3) deposited by the sole histone methyltransferase, Clr4 (Grewal & Jia, 2007). <u>Cl</u>r4 associates with <u>R</u>ik1 to form a <u>c</u>omplex (CLRC) that contains the ubiquitin E3 ligase Cul4-Rik1^Raf1/2^. This complex is essential for H3K9me establishment, but its ubiquitylation substrate remains unknown (Fig. 1A) (Hong *et al*, 2005; Horn *et al*, 2005; Jia *et al*, 2005; Li *et al*, 2005; Thon *et al*, 2005). H3K9me is recognized by members of the heterochromatin protein 1 (HP1) family, Swi6 and Chp2. HP1 proteins mediate spreading and the assembly into a heterochromatic platform by recruiting other factors, like SHREC (<u>S</u>nf2-like/<u>H</u>DAC-containing <u>re</u>pressor <u>c</u>omplex) (Motamedi *et al*, 2008; Sugiyama *et al*, 2007), which consists of Mit1, Clr3, Clr2, and Clr1 (Fig. 3A) (Sugiyama *et al*, 2007; Job *et al*, 2016). In addition, SHREC is targeted to subtelomeres via its interaction with Ccq1 (Moser *et al*, 2015; Sugiyama *et al*, 2007). Ccq1 also recruits CLRC and promotes silencing of reporter genes inserted next to telomeric repeats (Moser *et al*, 2015; Wang *et al*, 2016). However, unlike Ccq1, heterochromatin factors are not required for telomere maintenance (Ekwall *et al*, 1996). Telomere-adjacent (TAS) and telomere-distal subtelomeres differ in their requirements for heterochromatin silencing (Greenwood & Cooper, 2012), and the heterochromatin structure of the former is largely unknown.

Here we describe a role of Ccq1 in nucleosome maintenance at telomere-adjacent sub-telomeres. We identified Ccq1 as a main interactor and recruiter of CLRC, being critical for H3K9me deposition near telomeres, in agreement with two previous reports (Wang *et al*, 2016; Zofall *et al*, 2016). However, we further demonstrate that the major role of Ccq1 in heterochromatic silencing arises from maintaining critical levels of nucleosomes at the subtelomeric TAS. This chromosomal region displays unusually low nucleosome abundance, which is independent of the proximity of the telomeric repeats but intrinsic to the DNA sequence of the TAS. Deletion of *ccq1*^*+*^ results in a near complete loss of histones at the TAS. This unexpected function of Ccq1 in nucleosome maintenance is mediated by CLRC and SHREC, which are recruited by Ccq1 and act epistatically with Ccq1 in heterochromatic silencing. Thus, in contrast to other heterochromatic domains, gene silencing at the TAS appears to be regulated at the level of nucleosome stability. Fur-thermore, we show that recombination in the absence of Ccq1 is promoted by homologous sequences present in the TAS, implying that these sequences are actively implicated in telomere maintenance by a mechanism resembling ALT (alternative telomere lengthening).

## RESULTS

### 1. CLRC promotes the shelterin-chromatin association

To identify physical interactors of CLRC, we used a proteomic approach employing an epitope fusion with Raf1, the putative receptor of the Cul4-Rik1 E3 ubiquitin ligase. LC-MS analysis retrieved all members of CLRC as well as the shelterin complex (Suppl. Fig. S1A-C; Suppl. Table S1). This association required the presence of the adaptor protein Rik1 and was confirmed by co-immunoprecipitation experiments for the shelterin subunits Ccq1 and Tpz1 (Suppl. Fig. S1D). We further validated the Rik1-dependent interaction between CLRC and shelterin by a reciprocal approach expressing Ccq1^HA^ in WT and *rik1*Δ cells (Fig. 1A-B). In the absence of Rik1, the interaction pattern of Ccq1 was significantly altered, resulting in the loss of the interaction with CLRC and most of the other Ccq1-associated proteins (except the SHREC subunit Clr3 and the histone chaperone Hip1; Fig. 1B, right panel). Interestingly, we found several new interactions with factors involved in protein folding, cytokinesis, and non-homologous end-joining (NHEJ). Together, our proteomics data show that the shelterin complex is a main interactor of CLRC, confirming two recent reports (Wang *et al*, 2016; Zofall *et al*, 2016). We further demonstrate that the interaction profile of Ccq1 is affected by the presence of CLRC, suggesting that this association confines the co-operation of shelterin with other factors

Although CLRC contains a ubiquitin ligase subcomplex (Cul4-Rik1^Raf^1^/Raf2^), our results show that CLRC does not regulate shelterin through ubiquitylation (Suppl. Fig. S1E-F). Ccq1 and other shelterin subunits are enriched at telomeric repeats and the telomere-adjacent TAS. We therefore tested whether CLRC regulates the abundance of Ccq1 on subtelomeres and analyzed several loci next to the telomeric repeats known to bind the shelterin complex (Fig. 1C; see also Suppl. Fig. S2A-B) (Harland *et al*, 2014; Sugiyama *et al*, 2007; Tomita & Cooper, 2008). Lack of Rik1 or Clr4 causes a significant decrease in the binding of Ccq1, particularly next to the telomeric repeats (Fig. 1C). Similarly, association of all other shelterin components with chromatin is reduced in *rik1Δ* cells (Fig. 1D), except for Pot1. Since the latter binds the ssDNA overhang of the telomeric 3’ end in the semi-open confirmation (Miyoshi *et al*, 2008), we speculate that binding to the G-tail overhang is less sensitive to perturbations caused by the absence of Rik1. CLRC promotes heterochromatin assembly through the recruitment of HP1, however, neither of the single mutants nor the *swi6Δ chp2Δ* double mutant show a decrease in Ccq1 binding (Fig. 1E and data not shown). This suggests that CLRC-mediated stable binding of shelterin is not a direct function of the heterochromatin structure. To study whether CLRC can promote Ccq1 binding to chromatin independently of telomeres, we expressed Ccq1^HA^ in a strain in which the TetR^off^-Clr4* fusion protein can be tethered via a *4xtetO* array to an ectopic chromatin site (Audergon *et al*, 2015). Ccq1 associates with this ectopic site in a manner that strictly depends on the expression and binding of TetR^off^-Clr4* to the *tetO* array (Fig. 1F). Thus, CLRC promotes shelterin association with chromatin in a ubiquitylation- and HP1-independent manner.

### 2. Ccq1 promotes nucleosome stability at native subtelomeres

In agreement with two previous reports (Wang *et al*, 2016; Zofall *et al*, 2016), we found that both Rik1 and Raf1 associate with telomere-adjacent chromatin in a Ccq1-dependent manner, indicating that shelterin also recruits CLRC to chromatin (Suppl. Fig. S3A-B). We also observed that H3K9me2 is strongly reduced in *ccq1Δ* cells at TAS (Fig. 2A-B, left panel, and Suppl. Fig. S3C). This H3K9me2 loss was also seen for the *m23::ura4*^*+*^ reporter gene placed next to telomeric repeats and for an intrachromosomal Taz1-binding site (*isl15*) known to recruit the shelterin complex (Fig. 2A-B, middle and right panel), confirming results of the previous studies (Zofall *et al*, 2016; Wang *et al*, 2016). Nonetheless, we noticed that the chromatin distribution of CLRC markedly differs from the subtelomeric H3K9me2 profile. Whereas Rik1 and Raf1 are mostly abundant at the telomeric end and decline with increasing distance, H3K9me2 is mostly enriched at the heterochromatic *tlh1*^*+*^ gene located 15 kb away of the telomeric repeats (compare Suppl. Fig. S3A-B with Fig. 2). We further observed that H3K9me2 levels at the telomere-proximal TAS are unusually low, even in WT cells, reaching barely 10% of the levels found at *tlh1*^*+*^ (Fig. 2B, left panel). This low H3K9me2 level at TAS contrasts the levels seen at *m23::ura4*^*+*^ and the heterochromatin island *isl15* (Fig. 2B; middle and right panel). Hence, we wondered whether TAS are restrictive for H3K9 methylation or whether this chromatin region has lower nucleosome occupancy. ChIP for total histone H3 revealed that WT cells have indeed lower nucleosome levels at the telomere-proximal TAS1 region (0.3-fold compared to euchromatin, EC) compared to the telomere-distal *tlh1*^*+*^ locus (1.25-fold relative to EC; Fig. 2C). In the absence of Ccq1, H3 levels are even further decreased (0.1-fold at TAS1 relative to EC; Fig. 2C), but this marked decrease was not seen for *tlh1*^*+*^ or other heterochromatic loci (*m23::ura4*^*+*^, *isl15*; Fig. 2C; middle and right panel). By performing several control experiments (Supplemental Fig. S3E-F), we excluded the possibility that the H3 loss at TAS is caused by progressive telomere shortening, a phenotype often seen for cells upon deletion of *ccq1*^+^ after continuous growth (Tomita & Cooper, 2008). When determining the relative decrease, i.e. WT:mutant ratios for H3 and H3K9me2, we found a strong correlation for the telomere-proximal TAS, suggesting that the loss of H3 largely explains the loss of H3K9me2 at this subtelomeric region (Fig. 2D). Nonetheless, H3K9me2 levels are still significantly lower than H3 levels even in WT cells; thus, additional mechanisms likely contribute to constrain this repressive mark at TAS. In conclusion, our findings demonstrate that the chromatin structure at native subtelomeres differs substantially from other heterochromatin domains and reveals an unexpected role of Ccq1 in nucleosome maintenance.

**Figure 2.**
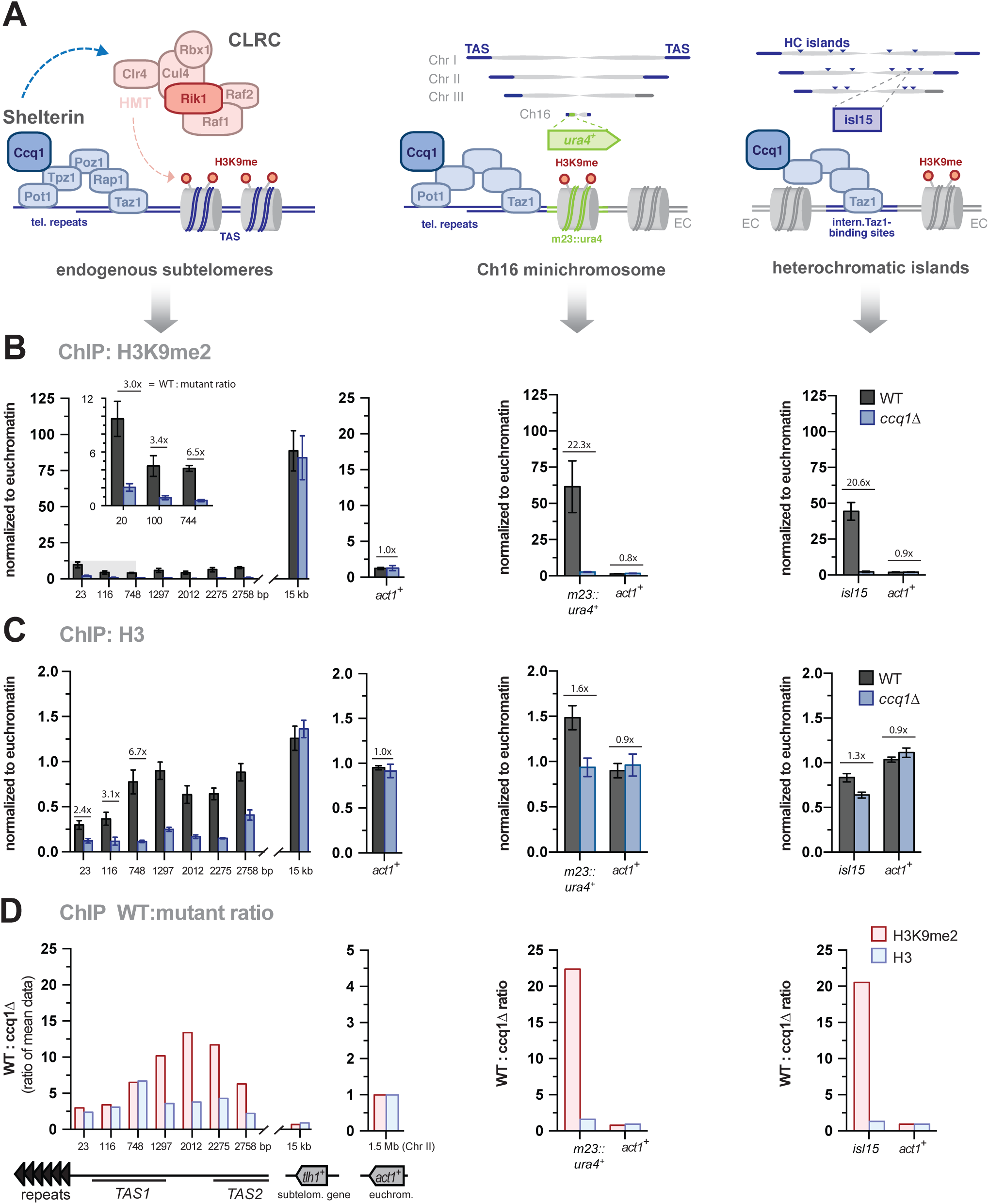
Ccq1 promotes nucleosome stability at native subtelomeres. **(A)** Left: Schemes of CLRC and the shelterin complex. Middle and right: Scheme depicting the position of the *ura4*^*+*^ reporter gene and heterochromatin islands relative to telomeric repeats and TAS regions; note that the mini-chromosome Ch16 does not contain TAS. **(B-C)** ChIP-qPCR analyses of H3K9me2 (B) and H3 (C) in WT and *ccq1 Δ* cells. ChIP data are normalized to input and to the average of three euchromatic loci (EC) as internal control (see methods), and represented as mean ± SEM from 3 independent experiments. **(D)** Fold change of WT over *ccq1Δ* for H3K9me2 and H3 levels (see methods).

**Figure 3.**
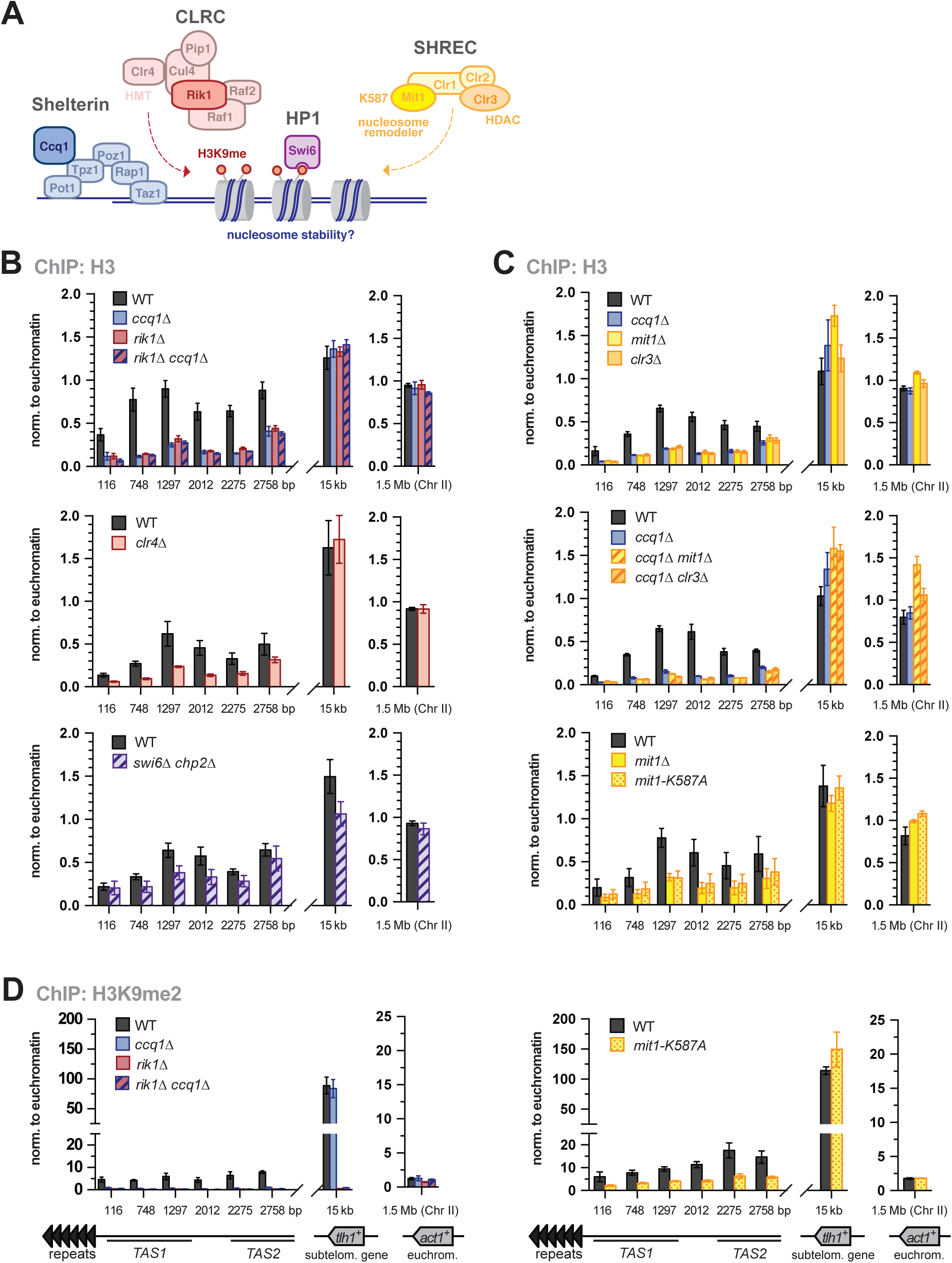
Ccq1 maintains nucleosome stability through recruiting CLRC and SHREC. **(A)** Scheme of shelterin, CLRC and SHREC complexes and Swi6^HP1^ bound to H3K9me2. **(B-D)** ChIP-qPCR analyses of H3 and H3K9me2 in WT and mutants as indicated. ChIP data are represented as mean ± SEM from 3 independent experiments (normalization as in Fig. 2B and C).

### 3. Ccq1 maintains nucleosome stability by recruiting CLRC and SHREC

Ccq1 may contribute to nucleosome stability through the recruitment of CLRC, which promotes H3K9 methylation and in turn prevents transcription-dependent histone turnover by RNA polymerase II (reviewed in (Workman, 2006)). To test whether CLRC acts downstream of Ccq1 in nucleosome maintenance, we analyzed H3 levels in mutants of CLRC. H3 levels are reduced to similar levels in single and double mutants of *rik1*^*+*^ and *ccq1*^*+*^, implying that the loss of CLRC alone is sufficient to produce this nucleosome loss phenotype (Fig. 3B, upper panel). Similar phenotypes were observed for *clr4Δ* cells and, albeit to lesser extent, for the *swi6*Δ *chp2*Δ double mutant (Fig. 3B, middle and lower panel). The latter is consistent with the fact that HP1 proteins provide a binding platform for CLRC (and SHREC, see below) and likely act redundantly with Ccq1 at the TAS. In agreement with the findings for *ccq1Δ,* the loss of H3 in mutants of CLRC is specific for TAS and not seen at *tlh1*^*+*^ (Fig. 3B).

Ccq1 was further shown to interact with the HDAC Clr3, which is part of the repressor complex SHREC that also contains the Snf2-like nucleosome remodeler Mit1 (Fig. 3A) (Creamer *et al*, 2014; Job *et al*, 2016; Moser *et al*, 2015; Sugiyama *et al*, 2007). Mit1 and Clr3 have been associated with nucleosome positioning and turnover and prevent the formation of nucleosome-depleted regions within heterochromatin (Aygün *et al*, 2013; Creamer *et al*, 2014; Sugiyama *et al*, 2007; Yamane *et al*, 2011). Similar to mutants of CLRC, we found that H3 is strongly decreased in *mit1Δ* or *clr3Δ* mutants at TAS, while the corresponding double mutants with *ccq1*Δ displayed an epistatic phenotype (Fig. 3C, upper and middle panel). To test whether the remodeler activity of Mit1 is required for nucleosome stability at native subtelomeres, we examined telomeric chromatin in a strain expressing an ATPase-dead mutant, *mit1-K587A* (Sugiyama *et al*, 2007). H3 levels in this point mutant are similar to the ones observed in *mit1Δ* (Fig. 3C, lower panel). This decrease in H3 at TAS is also reflected by reduced H3K9me2 in the *mit1-K587A* mutant, largely recapitulating the phenotype of *ccq1Δ* but differing from *rik1Δ,* which lacks H3K9me2 at *tlh1*^*+*^ (Fig. 3D). This finding further corroborates the notion that low levels of nucleosomes, or their improper positioning, result in fragile chromatin, precluding the establishment of H3K9me2 at TAS. In conclusion, Ccq1 maintains nucleosomes at telomere-adjacent subtelomeres through the recruitment of CLRC and SHREC.

### 4. The subtelomeric DNA sequence causes nucleosome instability

It has been proposed that subtelomeric sequences in *S. pombe* contain low affinity nucleosome binding sites forming nucleosome-free regions (Creamer *et al*, 2014; Garcia *et al*, 2010; Yamane *et al*, 2011; Sugiyama *et al*, 2007); however, these studies focused primarily on heterochromatin regions excluding the TAS regions. Employing ChIP-seq for H3, we find that telomere-proximal H3 levels in WT cells are low across the first 10 kb including the TAS, but rise with increasing distance from the telomeric ends (Fig. 4A). TAS2 and TAS3 have above-average A/T content (i.e. more than 70%) and a high percentage of exclusive A/T tracks of 5-mers and longer (Fig. 4B). Such DNA properties are absent at *tlh1*^*+*^ and further telomere-distal subtelomeric regions, which all display high levels of H3. Using an *in silico* nucleosome prediction algorithm (Xi *et al*, 2010), we found a strong correlation between our experimental H3 data and predicted nucleosome occupancy (Fig. 4C). Moreover, when analyzing mutants lacking Ccq1, Rik1 or Mit1, we found H3 levels to be further reduced throughout the TAS but to be mostly unaffected at the *tlh1*^*+*^ locus (Supplemental Fig. S4).

**Figure 4.**
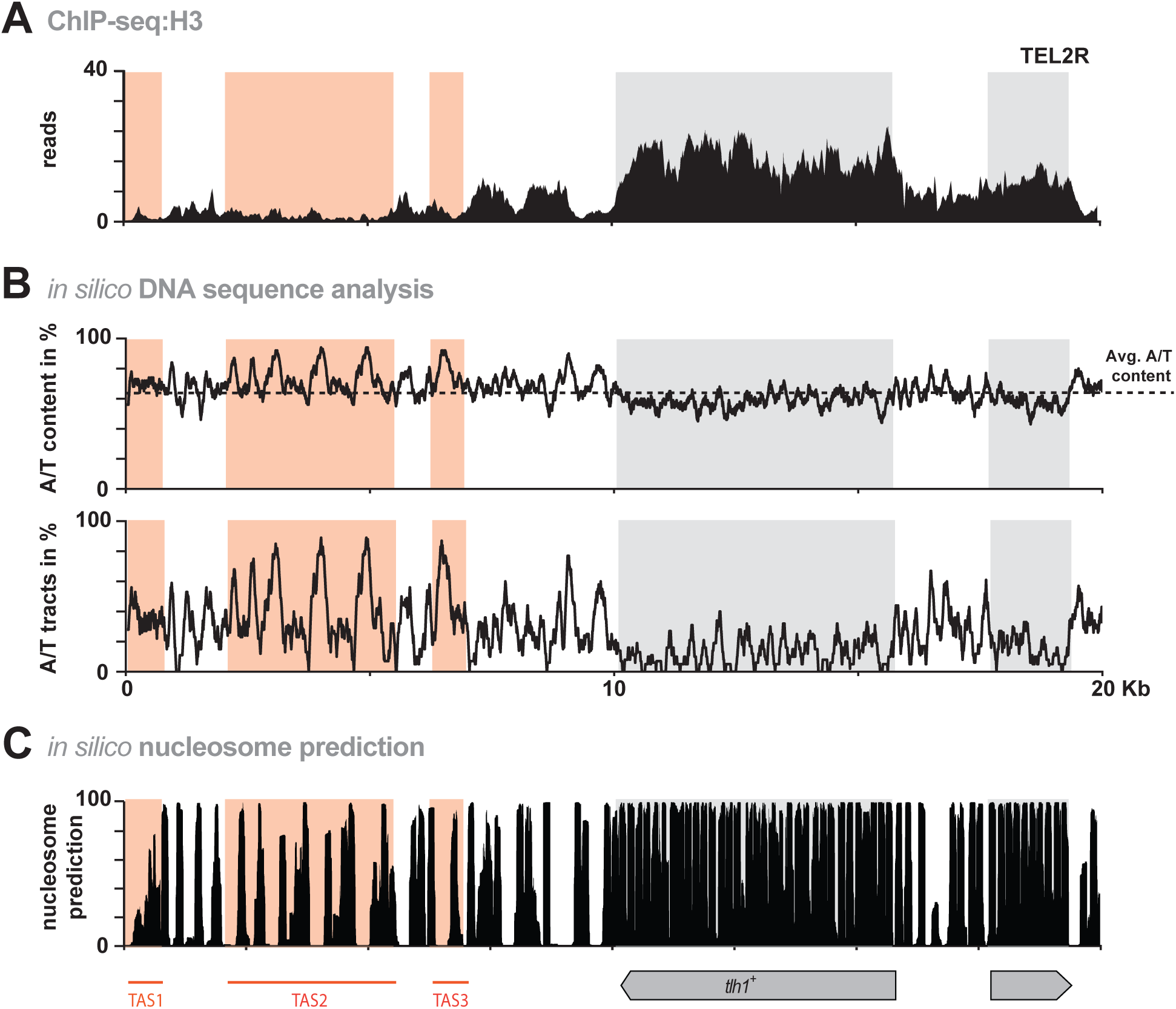
Nucleosome abundance is low throughout the telomere-proximal chromosomal region. **(A)** ChIP-seq reads mapped to TEL2R (shown in reverse orientation for consistency). **(B)** *In silico* analyses of TEL2R DNA sequence. Black line in top panel shows A/T content. Dotted line represents average A/T content in S. pombe (www.pombase.org) Bottom panel shows the percentage of poly(dA:dT) tracts (defined as a sequence of 5 or more nucleotides consisting only of A or T). Red and grey shaded areas show TAS regions and subtelomeric genes, respectively. **(C)** Nucleosome prediction for TEL2R using prediction algorithm (Xi *et al,* 2010).

To examine whether the low nucleosome occupancy at the TAS is an intrinsic feature of the underlying DNA sequence, we integrated a 790 bp fragment comprising a partial sequence of TAS1 (115-905 bp) into the intrachromosomal *leu1*^*+*^ locus using a plasmid-insertion strategy. Since TAS sequences are present at most of the chromosomal arms, distinguishing between subtelomeric and ectopic TAS sequences is difficult. We therefore integrated this TAS fragment in a strain that lacks all subtelomeric homologous (SH) sequences, including the TAS on chromosome I + II and the left arm on chromosome III (Tashiro *et al*, 2017). ChIP-qPCR revealed that H3 levels are low at the ectopic TAS1 fragment, recapitulating the H3 levels and distribution of the endogenous TAS1 region (Fig. 5A). Notably, H3 levels were further reduced at the ectopic TAS fragment when *ccq1*^*+*^ was deleted in this strain, similar to the endogenous TAS, suggesting that shelterin can bind to this fragment even in the absence of telomeric repeats (Fig. 5A). Together, these results indicate that the DNA sequence of this TAS fragment is sufficient to mediate nucleosome instability and regulation by Ccq1.

**Figure 5.**
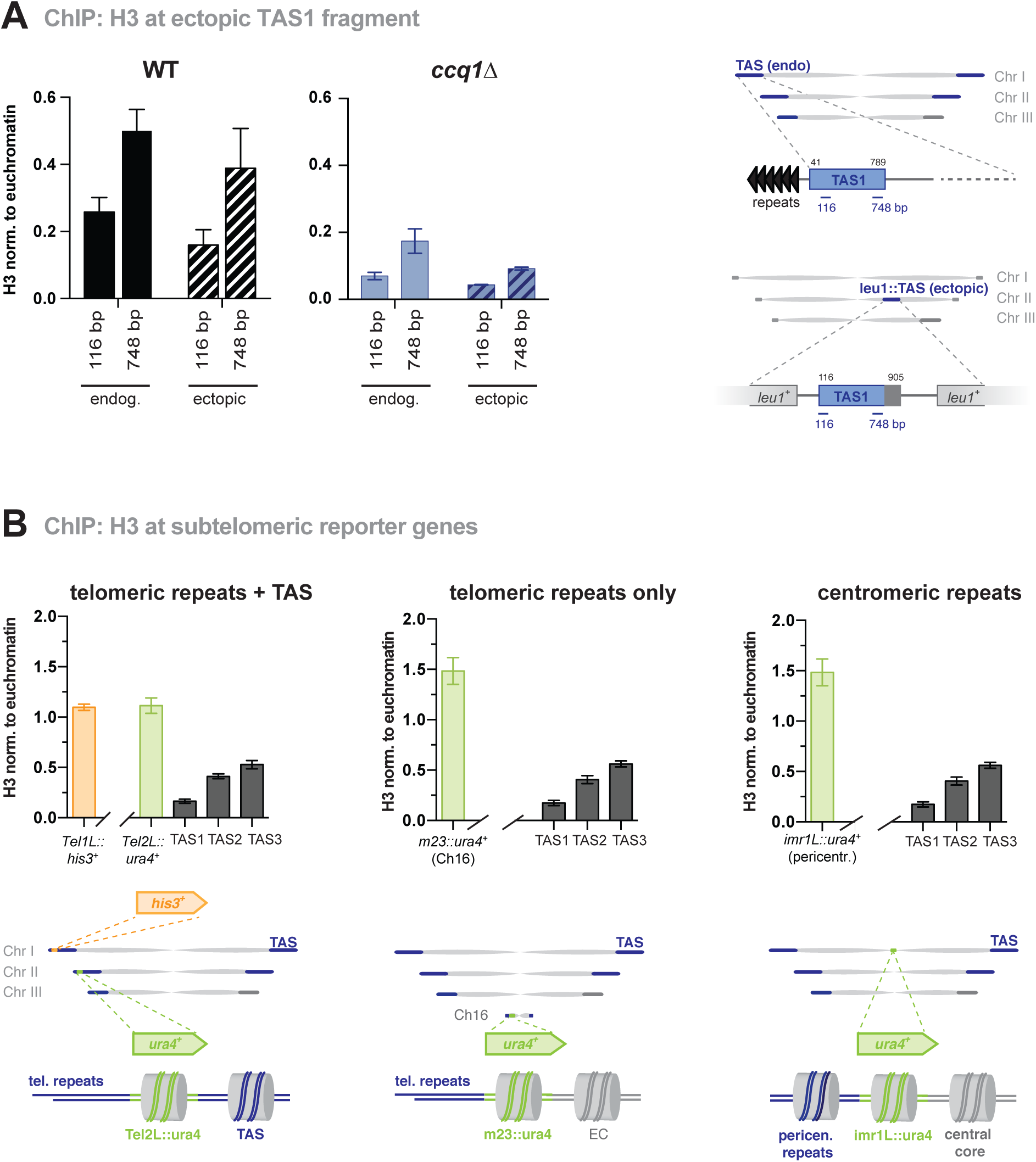
Ectopic insertion of TAS fragment is sufficient to cause nucleosome instability. **(A)** ChIP-qPCR analysis of H3 at endogenous TAS and a fragment spanning the TAS region from 115-905 bp (relative to telomeric repeat) inserted into the *leu1*^*+*^ locus (see scheme). The TAS fragment was inserted into a strain that lacks endogenous TAS (see text) (Tashiro et al, 2017). Shown are ChIP analyses for WT (left) and *ccq1Δ* (right; note different the scale of the y-axis). Data are represented as mean ± SEM from 9-10 independent experiments (normalization as in Fig. 2C) except for ectopic TAS in *ccq1Δ* strain where n =3. **(B)** ChIP-qPCR analysis of H3 at reporter genes (*ura4*^*+*^ and *his3*^*+*^) at various chromosomal locations (see schemes) in WT cells. TAS1, TAS2, and TAS3 correspond to position 116 bp, 2851 bp and 6291 bp (relative to telomeric repeats), respectively. Data are represented as mean ± SEM from 3 independent experiments (normalization as in Fig. 2C).

To investigate whether the proximity to telomeres also contributes to nucleosome instability, we examined H3 levels at the *ura4*^*+*^ gene inserted between the telomeric repeats and TAS on the left arm of chromosome II (*Tel2L::ura4*^*+*^; scheme see Fig. 5B, left) (Nimmo *et al*, 1998). In analogy to *ura4*^*+*^ inserted at the *m23* locus on the Ch16 minichromosome (Nimmo *et al*, 1994), *Tel2L::ura4*^*+*^ is marked by H3K9me2 and efficiently silenced (Nimmo *et al*, 1998). Yet in contrast to TAS, *Tel2L::ura4*^*+*^ and *m23::ura4*^*+*^ display high H3 levels in WT cells, which are comparable to H3 levels of *ura4*^*+*^ inserted at the pericentromeric *imr1L* locus (Fig. 5B). These high H3 levels at *m23::ura4*^*+*^ and *Tel2L::ura4*^*+*^ are mostly maintained in *ccq1Δ* cells (Fig. 2C and Suppl. Fig. S5A). We made similar observations for the *Tel1L::his3*^*+*^ reporter gene present on the left arm on chromosome I (Fig. 5B). Based on these findings, we conclude that the presence of the telomeric repeats is not sufficient to cause nucleosome instability. We also tested whether the shelterin complex competes with nucleosomes for DNA binding by analyzing H3 levels in cells lacking Taz1, which recruits shelterin to dsDNA (Cooper *et al*, 1997; Miyoshi *et al*, 2008). However, deleting *taz1*^*+*^ resulted in reduced H3 levels similar to *ccq1Δ* or *mit1Δ.* Consistent with this finding, nucleosome levels were not recovered in *taz1Δ* double mutants lacking Ccq1 or Mit1 (Suppl. Fig. S5B), indicating that removing shelterin from subtelomeres is not sufficient to restore nucleosome levels, neither to levels seen WT cells nor to levels found at other genomic regions. Together, these findings demonstrate that the specific chromatin context of the underlying DNA sequences, or the DNA sequence itself, but not the chromosomal positioning, determines nucleosome stability at TAS.

### 5. Subtelomeric nucleosome stability is linked to heterochromatic silencing

To understand the impact of nucleosome stability on telomere function, we analyzed sub-telomeric silencing by examining the expression of endogenous subtelomeric non-coding RNAs (here dubbed as *TERRA,* telomeric repeat-containing non-coding RNA, for simplicity) from telo-mere-adjacent TAS regions (Suppl. Fig. S6A) (Bah *et al*, 2012; Greenwood & Cooper, 2012). Mutants of *ccq1* and *mit1* cause comparable levels of *TERRA* derepression and display an epistatic genetic interaction (Fig. 6A, left panel), similar to that observed for nucleosome stability. Loss of Clr3 and Rik1 resulted in a slightly larger increase of *TERRA* and showed partially additive effects with *ccq1Δ*, suggesting that they can also act independently in *TERRA* silencing (Fig. 6A, left and right panels). Similar observations were made for the binding of RNA polymerase II to chromatin regions associated with *TERRA* transcription (Suppl. Fig. S6B). However, in contrast to the mostly non-additive derepression of *TERRA*, we found that *ccq1Δ* double mutants also deficient in SHREC or CLRC display strong synthetic silencing defects at *tlh1*^+^ (Fig. 6B). This led us to conclude that Ccq1 contributes to silencing by different mechanisms: at telomere-adjacent chromatin, Ccq1 acts to maintain nucleosome stability mainly through the combined actions of SHREC and CLRC, whereas at *tlh1*^+^, Ccq1 has a different function that is independent of SHREC and CLRC.

**Figure 6.**
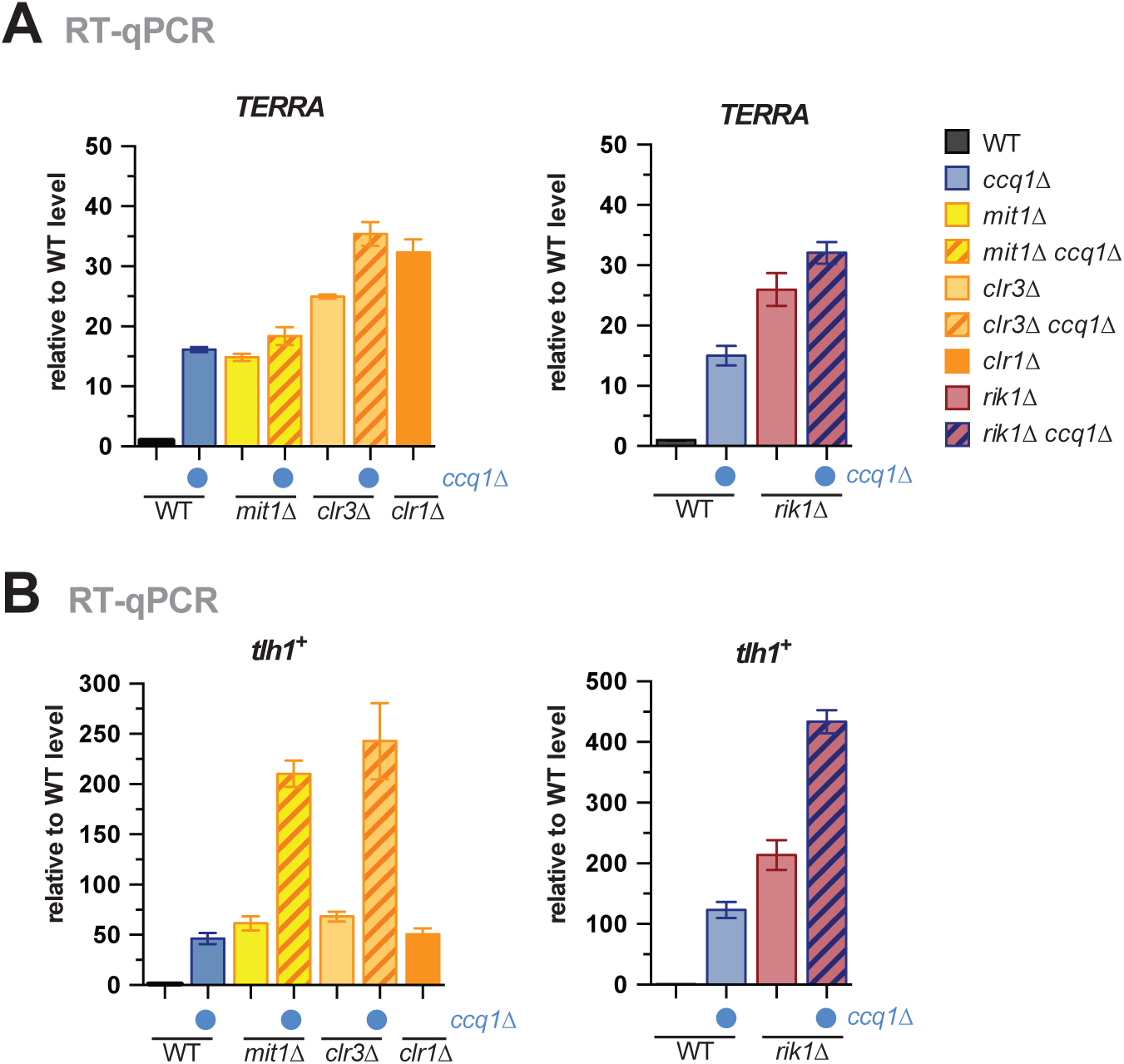
Nucleosome stability at subtelomeres is linked to heterochromatic silencing. **(A, B)** RT-qPCR analysis of transcript levels of *TERRA* (A) and *tlh1*^*+*^ (B) in WT strain and mutants as indicated (double mutants with *ccq1Δ* are indicated by blue dot). RT-qPCR data is represented as mean ± SEM from 3 independent experiments and shown relative to WT level (see methods).

### 6. TAS promote subtelomeric DNA recombination in the absence of Ccq1

Our findings indicate that the intrinsic properties of the TAS sequence impede nucleosome binding, raising the question about the benefit of maintaining these sequences at subtelomeres. Deletion of Ccq1 was previously shown to cause genomic rearrangements of subtelomeres, which likely contribute to telomere maintenance in the absence of telomerase (Tomita & Cooper, 2008). Thus, we wondered whether TAS are critical for this hyper-recombinogenic phenotype of *ccq1Δ*. While we did not observe changes in the overall abundance of TAS under our experimental conditions (Suppl. Fig. S3F), recombination events between individual subtelomeres (e.g. gene conversion or break-induced replication) would be difficult to detect due to the identical DNA sequences on most chromosomal arms. To overcome this technical challenge, we used reporter genes inserted into telomeric regions that served as unique barcodes. As such, we used *ura4*^*+*^ placed next to telomeric repeats on the minichromosome Ch16 (*m23::ura4*^*+*^), which lacks any TAS sequences (Nimmo *et al*, 1994), and compared its stability with *his3*^*+*^ and *ura4*^*+*^ inserted between the endogenous telomeric repeats and TAS1 (*Tel1L::his3*^*+*^ *Tel2L::ura4*^*+*^) (Nimmo *et al*, 1998). Cultures of WT cells untreated or freshly disrupted for *ccq1*^*+*^ were grown continuously for about 40 generations, and genomic copy numbers of the reporter gene and TAS were assessed every six to seven generations (see scheme, Fig. 7A).

**Figure 7.**
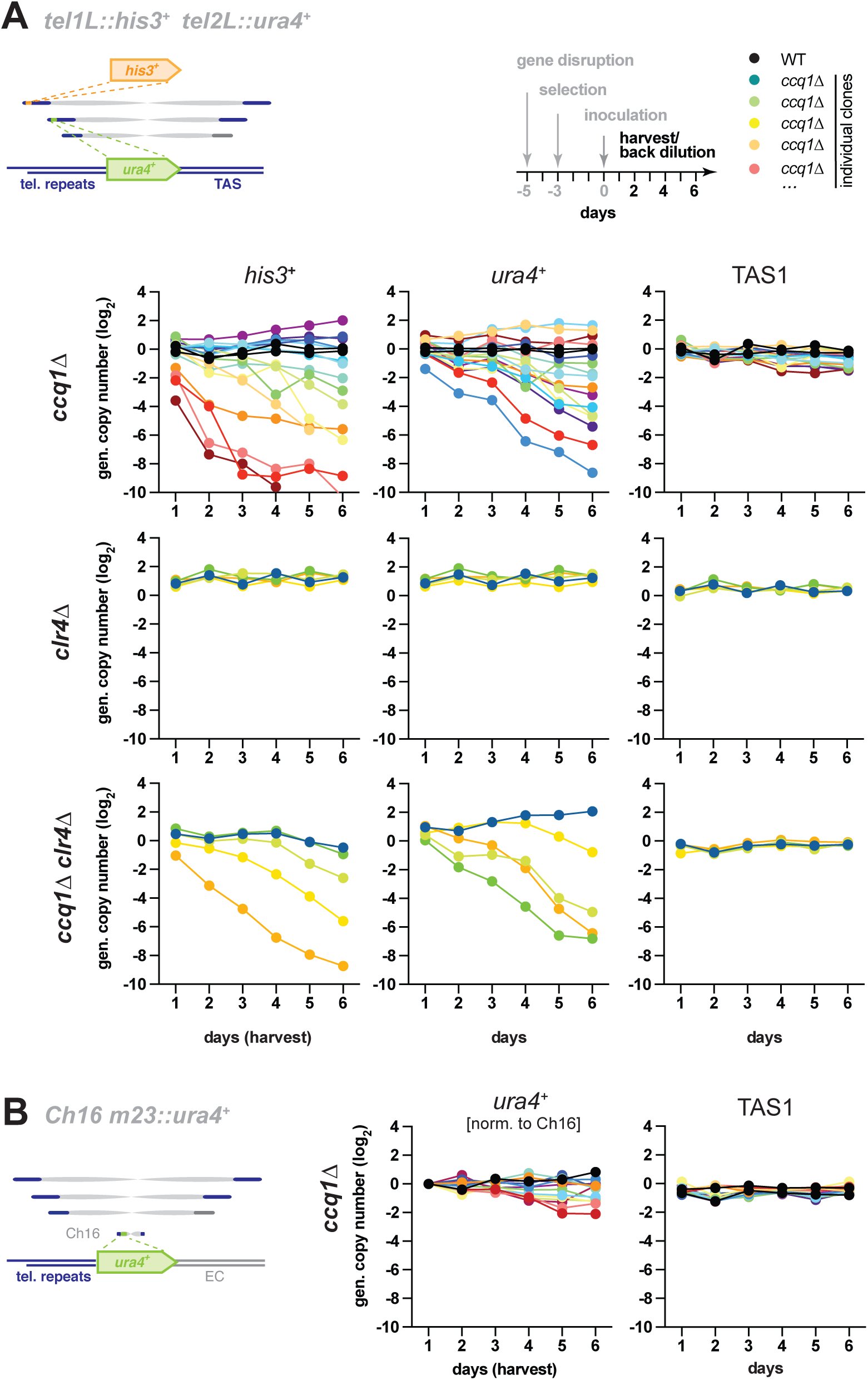
Subtelomeric DNA promotes recombination in the absence of Ccq1. Genomic copy number of *his3*^*+*^, *ura4*^*+*^ and TAS1 in indicated strains harboring the reporter genes *tel1L::his3*^*+*^ and *tel2L::ura4*^*+*^. Cultures from individual WT and freshly generated knockout clones (*ccq1Δ,* n = 16;, *clr4Δ,* n = 5; *ccq1Δ clr4Δ, n = 5)* were pre-grown on selective media for several days and inoculated at day 0 to grow in liquid media with regular back-dilution (every 24 hours, approximately 7 generations). Samples were taken at indicated harvest times, and relative copy numbers of genomic regions were assessed by qPCR (normalization against intrachromosomal loci). Black and rainbow color lines indicate WT strains and individual *ccq1*^*+*^ deletion mutants (clones), respectively. **(B)** qPCR analysis as in (A) but with strains harboring the minichromosome *Ch16 m23::ura4*^*+*^ (*ccq1Δ,* n = 14). Since Ch16 is unstable in *ccq1Δ* cells (see Suppl. Fig S8A-B and Motwani *et al*, 2010), *m23::ura4*^*+*^ has been normalized to minichromosome levels (details on normalization, see methods). Note that Ch16 does not harbor TAS sequences.

In contrast to WT cells, we found that both *Tel1L::his3*^*+*^ and *Tel2L::ura4*^*+*^ were highly unstable in *ccq1Δ* cells, resulting in either additional copies or their gradual loss, but largely independently of each other (Fig. 7A and Supplemental Fig. S7). Moreover, whereas most isolates displayed a partial or near complete loss of the reporter gene in the culture, those with a copy number gain showed a maximum of 4-5 copies of either *his3*^*+*^ or *ura4*^*+*^, suggesting that most TAS-containing subtelomeres acquired an additional copy of the reporter gene in these cultures. We did not observe major changes for the TAS1-3 regions (Fig. 7A and data not shown), suggesting that the rearrangements of subtelomeric *his3*^*+*^ and *ura4*^*+*^ were largely caused by recombination but not telomere shortening. In contrast to the TAS-flanked reporters, we found that the *m23::ura4*^*+*^ reporter gene is relatively stable in *ccq1Δ* cells (Fig. 7B and Supp. Fig. S8), particularly when taking into account the general instability of the Ch16 minichromosome in this mutant reported previously (Motwani *et al*, 2010).

To dissect whether TAS-mediated genome instability is linked to nucleosome loss or relates to other functions of Ccq1, we repeated these experiments in strains deficient in CLRC or SHREC. Examining reporter gene stability in single mutants of *clr4*^*+*^ (Fig. 7A, middle panel) or *rik1*^*+*^, *mit1*^*+*^ and *clr3*^*+*^ (Suppl. Fig. S7A) revealed no significant difference to WT cells. This implies that failed recruitment of CLRC and SHREC, and as consequence nucleosome loss, is not sufficient or involved to cause this phenotype. We further tested whether CLRC is required for recombination, in analogy to HAATI (heterochromatin amplification-mediated and telomerase-independent), an alternative telomere maintenance mechanism (Jain *et al*, 2010). Deleting *clr4*^*+*^ did not prevent the hyper-recombinogenic phenotype of *ccq1Δ* (Fig. 7A, lower panel), indicating that the requirements for HR involving DNA sequences present within TAS are different from HAATI (which predominantly involves rDNA). In conclusion, our findings imply that subtelomeric sequences are intrinsically prone to recombination, which is prevented by Ccq1 through a mechanism independent of its role in nucleosome maintenance.

## DISCUSSION

Here, we demonstrate that the subtelomeric TAS comprise a unique chromatin structure that is controlled by the shelterin subunit Ccq1. This chromosomal region displays very low nucleosome abundance, which is sequence-dependent but position-independent. Neither transcription, nor the presence of telomeric repeats, nor competition with shelterin explains these low levels (Fig. 5-6 and Supp. Fig. S5). Rather, the presence of exclusive poly(dA:dT) tracts strongly correlates with the low nucleosome occupancy (Fig. 4), and integration of a TAS fragment at an ectopic locus is sufficient to cause nucleosome instability (Fig. 5). Consistently, the role of Ccq1 in nucleosome stability, mediated by CLRC and SHREC, is strictly confined to TAS but not found at telomere-distal subtelomeres, which have normal nucleosome abundance. Interestingly, those telomere-distal regions still display Ccq1-dependent silencing, yet in a CLRC- and SHREC-independent manner, suggesting that Ccq1 contributes to heterochromatin maintenance at distinct regions by different mechanisms. Hence, it is an attractive hypothesis that Ccq1’s property to interact with CLRC and SHREC has co-evolved as a ‘built-in function’ with the appearance of subtelomeric nu-cleosome-refractory sites to accommodate the need of maintaining appropriate histone levels.

The low nucleosome occupancy at TAS may result from less well-positioned nucleosomes. This is in agreement with a recent study that describes a non-canonical chromatin structure at chromosomal ends consisting of irregularly spaced nucleosomes and telomere-binding proteins (‘telosomes’) (Greenwood *et al*, 2018). However, while this structure appears to be restricted to the ∼1.5 kb telomere-proximal region and requires the presence of chromosomal ends, we find that the low nucleosome occupancy and instability at TAS extends to chromosomal regions more distal (Fig. 4) and is independent of telomeric repeats (Fig. 5). This suggests that TAS act autonomously in controlling nucleosome binding, which appears to be encoded in the DNA sequence. The A/T-rich sequence may disfavor nucleosome binding (Creamer *et al*, 2014) or TAS may recruit in a se-quence-dependent manner other factors that compete with nucleosome binding. For example, Taz1 binds DNA beyond the telomeric repeats (Kanoh *et al*, 2005), which may be facilitated by low nucleosome occupancy. However, deleting *taz1*^*+*^ is not sufficient to restore nucleosome levels (Suppl. Fig. S5), suggesting that additional or other mechanisms are involved. We tested a potential role by the <u>R</u>emodels <u>S</u>tructure of <u>C</u>hromatin complex (RSC), which promotes nucleosome eviction and has been proposed to antagonize Mit1 (Creamer *et al*, 2014; Garcia *et al*, 2010). While TAS2 and TAS3 displayed a modest H3 increase in a temperature-sensitive mutant of RSC, this increase was even stronger at euchromatic genes, and overall nucleosome occupancy remained low at TAS compared to other genomic regions (Suppl. Fig. S5). Hence, we conclude that RSC does not play a specific role in the nucleosome instability at TAS. Interestingly, origins of replication correlate with low nucleosome occupancy (Lantermann *et al*, 2010) and the origin recognition complex (ORC) binds adenine stretches via its Orc4 subunit, which contains several AT-hook subdomains (Chuang & Kelly, 1999; Okuno *et al*, 1999). Thus, ORC may be a potential factor that competes with nucleosomes for binding to TAS.

Since nucleosomes are the major obstacle for transcribing RNA polymerase II, the absence of Ccq1, or its downstream partners CLRC and SHREC, likely have a direct consequence on gene expression. We find that the defect in maintaining H3 levels in these mutants correlates with the derepression of ncRNAs expressed from TAS (Fig. 6). The role of Ccq1 in repressing transcription could be directly mediated through the nucleosome remodeler activity of Mit1, which may help keeping nucleosomes at refractory binding sites, in agreement with previous reports (Creamer *et al*, 2014; Garcia *et al*, 2010; Yamane *et al*, 2011; Sugiyama *et al*, 2007). The activities of Clr3 and CLRC may additionally contribute to nucleosome stability through preventing open chromatin structure and access to RNA polymerase II, which would be consistent with the role of Clr3 in restricting histone turnover at other heterochromatic domains (Aygün *et al*, 2013). However, even when fully derepressed, those other heterochromatin domains maintain high nucleosome levels, whereas perturbing any of these mechanisms at TAS causes a substantial decrease of H3 with nearly a complete loss at some sites (Fig. 3). Thus, we conclude that transcription alone is not sufficient to cause the nucleosome loss at TAS, underscoring the fragile nature of this subtelomeric region, which makes it necessary to provide multiple mechanisms to maintain a critical level of nucleosomes and repression of *TERRA* (see model, Fig. 8A). On the other hand, since telomere elongation is promoted by the expression of *TERRA* (Moravec *et al*, 2016), the metastable state of nucleosomes may also serve as a cue to dynamically control telomere length. This is further in line with the negative regulation of telomere elongation by SHREC, which in addition displaces telomerase from shelterin (Armstrong *et al*, 2017).

**Figure 8.**
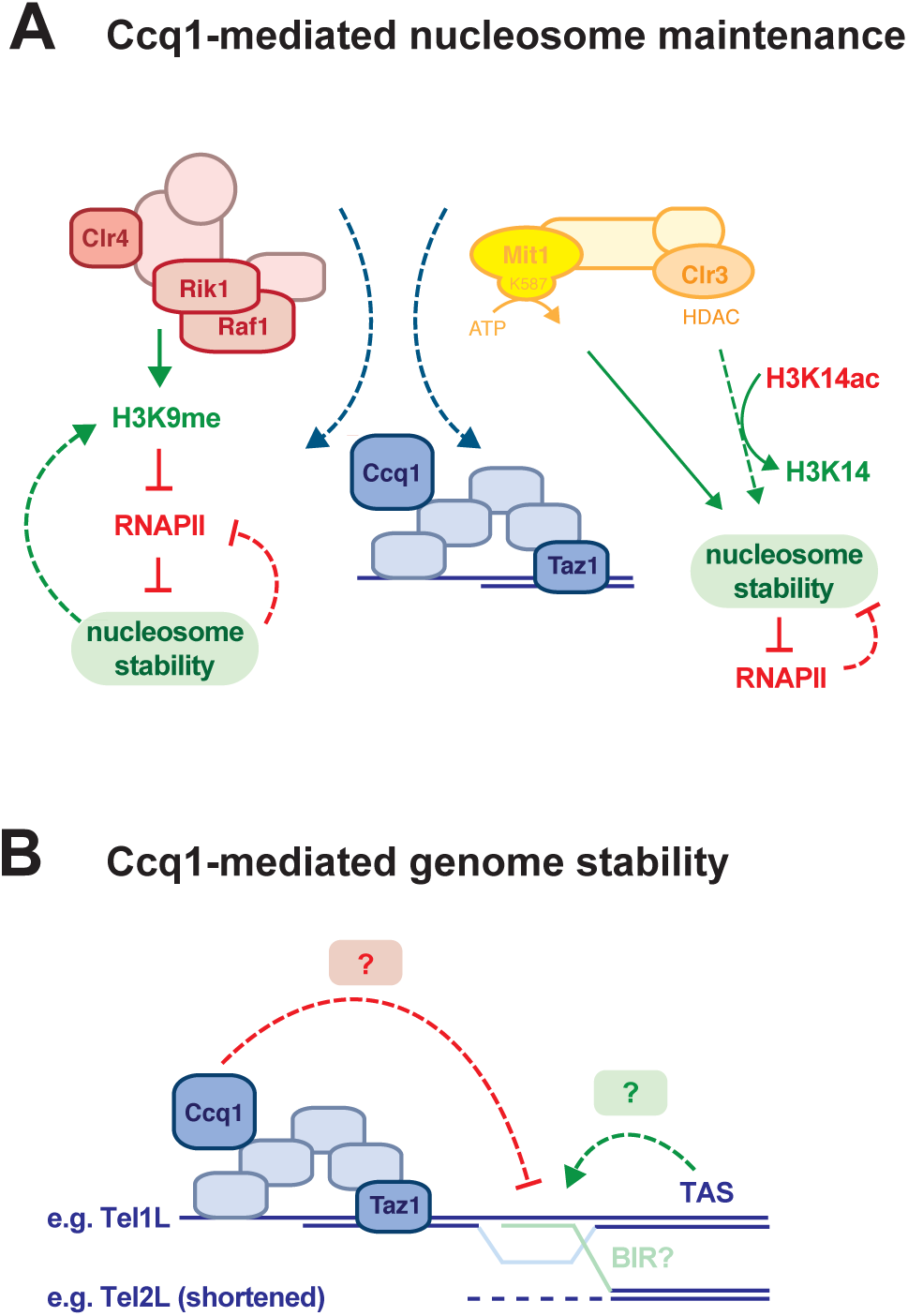
Working models for Ccq1-mediated nucleosome stability and telomere maintenance. **(A)** Ccq1 recruits CLRC and SHREC to subtelomeric chromatin, which counteracts nucleosome instability conferred by the DNA sequence of TAS. CLRC may contribute to nucleosome stability indirectly by depositing H3K9me, which prevents access to RNAPII and, thus, nucleosome eviction through transcription. Conversely, stabilized nucleosomes will contribute to maintaining critical levels of H3K9me at TAS. SHREC may contribute to nucleosome stability directly through nucleosome positioning via its ATP-dependent remodeler activity conferred by Mit1; indirectly through histone deacetylation via the HDAC moiety Clr3, thereby preventing open chromatin structure and access to RNAPII. **(B)** TAS or other DNA sequences may represent fragile sites causing DNA breaks through replication fork collapse or the recruitment of unknown factors causing genome instability. Ccq1 may prevent genomic instability by binding to subtelomeric chromatin, thereby stabilizing fragile sites and/or competing with the binding of destabilizing factors. Homologous sequences present in the TAS may promote recombination by BIR (break-induced replication) or similar mechanisms.

Taz1 and other shelterin proteins are known to be critical for H3K9 methylation (Kanoh *et al*, 2005; Tadeo *et al*, 2013; Zofall *et al*, 2016; Wang *et al*, 2016). The recruitment of CLRC by Ccq1 has therefore been proposed to be essential for robust heterochromatin assembly and the initiation of heterochromatin spreading from telomeric repeats to telomere-distal subtelomeres (Wang *et al*, 2016). However, our results suggest that the decrease in H3K9me2 in *ccq1Δ*, at least at telomere-adjacent subtelomeres, is less a cause but rather a consequence of nucleosome loss. This becomes very obvious in cells lacking Mit1, which results in a similar loss of H3K9me2 at TAS yet does not affect H3K9me2 maintenance at other heterochromatin domains (Fig. 3), consistent with previous findings (Creamer *et al*, 2014). Furthermore, TAS recruitment of CLRC by Ccq1 results only in low H3K9me levels in WT cells (Fig. 2), raising the question of whether these low H3K9me levels are sufficient to promote HP1-dependent spreading. These apparent discrepancies between our work and previous studies can likely be explained by H3K9 methylation and silencing having been primarily examined in the context of reporter genes (Kanoh *et al*, 2005; Tadeo *et al*, 2013; Wang *et al*, 2016). In contrast to native subtelomeres, these euchromatic genes exhibit normal nucleosome abundance and allow the establishment of high levels of H3K9me, even when placed next to telomeric repeats (Fig. 2). We made similar observations for nucleosome occupancy at intrachromosomal Taz1-binding sites (Fig. 2), which promote Ccq1-dependent assembly of facultative heterochromatin (Zofall *et al*, 2016). Thus, Ccq1 appears to fulfill distinct functions, promoting nucleosome maintenance and heterochromatin establishment, largely depending on the chromatin environment. From this we predict that heterochromatin at TAS is primarily controlled by nucleosome stability, in contrast to other heterochromatin regions that display high levels of H3K9me and depend on HP1-mediated spreading. Finally, the physical interaction between Ccq1 and CLRC may also impact other processes. In the absence of telomerase, linear chromosomes can be maintained through HAATI, which depends on Ccq1 and heterochromatic factors (Jain *et al*, 2010; Begnis *et al*, 2018). It has been proposed that Ccq1 could be recruited to heterochromatin via its interaction with Clr3 (Jain *et al*, 2010). Since we find that the association of Ccq1 with telomeres or an ectopic intrachromosomal locus is also controlled by CLRC (Fig. 1), this may suggest that CLRC also plays an active role in shelterin regulation and HAATI recruitment.

The persistent challenge of maintaining proper nucleosome levels raises the question of what is the advantage of keeping these nucleosome-binding refractory sequences at subtelomeres. Whereas SH sequences, which include TAS, are dispensable for mitotic and meiotic growth, their relevance for telomere maintenance becomes evident when Trt1 (the catalytic subunit of telomerase) is absent. Under this condition, loss of SH sequences shifts the balance among *trt1Δ* survivors from intra-to interchromosomal fusions absent (Tashiro *et al*, 2017). The conclusion that has been drawn from this observation is that the SH sequences promote circularization upon telomere erosion. However, in contrast to *trt1Δ* cells, early *ccq1Δ* mutants do not circularize, but instead maintain linear chromosomes, via gross genomic rearrangements, suggesting an additional function of subtelomeres. This pathway involves ATR (<u>A</u>taxia telangiectasia and <u>R</u>ad3 related)-dependent checkpoint activation and an unusual HR mechanism that depends on Rad51 but not Rad52 (Tomita & Cooper, 2008). We confirmed this hyper-recombinogenic phenotype of *ccq1Δ* and demonstrated that recombination involves DNA sequence present within TAS, suggesting that they are directly involved in the telomere maintenance mechanism that becomes active upon loss of Ccq1. As we observe that donor sequences often get amplified in *ccq1Δ* resulting in loss of heterozygosity, we hypothesize that recombination occurs through break-induced replication (BIR), resembling other telomere maintenance mechanisms in *S. cerevisiae* (McEachern & Haber, 2006) or in human tumor cells commonly referred to as ALT (alternative lengthening of telomeres) (Apte & Cooper, 2017).

Several mechanisms could contribute to the genomic instability at TAS. Formally, we cannot exclude the possibility that any homologous donor sequence would be sufficient to sustain a high recombination rate at subtelomeres. However, while the presence of homologous donor sequences likely affects the mode of repair (HR versus NHEJ), DNA damage at those hyper-recombinogenic sites may be due to additional intrinsic properties of the chromatin region. Thus, we favor the hypothesis that the genomic instability is specifically caused by the DNA sequence of the TAS (see model, Fig. 8B). For instance, recombination may be facilitated by the sequence properties of TAS (e.g. poly(dA:dT) tracts), which may be more prone to double-strand breaks (DSBs) (Schwartz *et al*, 2006). This idea would be consistent with our observation that TAS trigger recombination of telomere-proximal reporter genes, implying that DSBs frequently occur telomere-distal of the reporters within subtelomeric sequences when Ccq1 is not present. A similar role has been proposed for intrachromosomal Taz1-dependent heterochromatin islands, which replicate late (a feature linked to fragile sites) and map to meiotic DSB sites (Zofall *et al*, 2016). In addition, the mouse homolog of Taz1, TRF1, was shown to bind to a common fragile site containing an internal telomeric repeat and to stabilize this locus in a sequence-but position-independent manner (Bosco & de Lange, 2012). Alternatively, sequence-specific elements within the TAS may mediate the recruitment of specific factors promoting HR, as seen in human ALT cells. Here, rearrangements of telomere repeats are driven by the nuclear hormone receptor (NHR) family members NR2C/F. These receptors recognize variant telomere repeat motifs and are thought to dimerize and ‘bridge’ the canonical and variant repeats, resulting in their clustering and recombination (Marzec *et al*, 2015). Finally, ncRNAs derived from the subtelomeric TAS, such as *TERRA*, could also contribute to genome stability through the formation of RNA:DNA hybrids, as discussed for ALT (Apte & Cooper, 2017). However, although we find that TERRA transcription is upregulated in *ccq1Δ* and mutants of CLRC (Fig. 2-4), the absence of CLRC or SHREC alone is not sufficient to induce recombination (Supplemental Fig. S7). This is consistent with the minor role of heterochromatin in inhibiting related telomere recombination pathways (Khair *et al*, 2010) and implies that other downstream functions controlled by Ccq1 are critical in this process.

Many species harbor subtelomeres with mosaic homologous sequence elements and there is increasing evidence that they play a role in telomere length control independent of telomerase, as seen in yeast, flies, and humans (Jain & Cooper, 2010). Our findings demonstrate that the TAS in *S. pombe* differ from other chromatin regions through their nucleosome instability and high recombinogenic potential. Further studies of the phenomena may provide crucial insights into the still opaque functions of subtelomeres and, specifically, how telomere maintenance mechanisms are encoded in the subtelomeric DNA sequence.

## MATERIAL AND METHODS

### Yeast techniques, plasmids and strains

Standard media and genome engineering methods were used (Fission Yeast. A Laboratory Manual. Hagan, Carr, Grallert and Nurse. Cold Spring Harbor Press. New York 2016). 5-FOA media contained 1 g/L 5′-fluoroorotic acid. EMM-leu media were used for growing strains harboring *pREP1* plasmids. PMG medium supplemented with 10 µM anhydrotetracycline (AHT) and 15 µM thiamine was used for experiments with 4xTetO strains expressing FLAG-TetR-Clr4*. The plasmid used for generating the UBA trap fusion strains (3xFLAG-DSK2) was described previously (Mark *et al*, 2013). The plasmids *pBS-3xFLAG:kanMX6, pFA6a–HA:kanMX4*, and *pREP-nmt1p-His-ubi-LEU2* were provided by M. Smolle (LMU Munich), A. Ladurner (LMU Munich), and T. Toda (Hiroshima University), respectively. The ectopic TAS strain was generated by amplifying 790bp of TAS1 region using genomic DNA from the strain PSB0017 as a template and oligonucleotides Sg3182 and Sg3183. The PCR product was cloned into the plasmid *pJK148*. For integration into the *leu1-32* locus, the resulting plasmid was linearized using *Nde*I (NEB R0111S) and transformed into *S. pombe* strain ST3479, that lack endogenous TAS (gift by J. Kanoh, Osaka University). Strains used in this study are listed in Supplemental Table S2.

### Large Scale Immunoprecipitation (IP)

Cell extracts were prepared with 20-30 g of cells. Liquid cultures were harvested by centrifugation, washed with STOP Buffer (NaCl 150 mM, EDTA 10 mM pH 7.4, 2 mg/ml Sodium Fluoride, 0.065 mg/ml Sodium Azide), resuspended in 3.5-5 mL of lysis buffer (25 mM HEPES-KOH, 150 mM KOAc, 10 mM MgCl_2_, 5 mM CaCl_2_, 20% Glycerol, 4 mM AEBSF [Sigma, 11585916001], 1 Roche Complete EDTA-free, protease inhibitor tablet [Roche, 05056489001] per 10 ml, and 0.01 mg/ml MG132 [Sigma, C2211]); cell droplets were flash-frozen by pipetting into liquid nitrogen. Frozen droplet cells were broken during 9 cycles (3 min ON/2 min OFF, power 9) with a freezer mill [SPEX, 6970EFM-50 mL adaptor]. Cells were diluted up to 0.6 g/ml and treated with 174 U/ml of Benzonase for 1h at 4°C. Cell debris was discarded (17530 g, 15 min at 4°C), and the soluble fraction was clarified by centrifuging at 40K for 45 min at 4°C. After a pre-clearing step with Dynabeads Prot G [Life Technologies, 10009D], the supernatant was incubated with the Dynabeads Prot G beads coupled to the respective antibody for 3-12 h. The bound material was washed two times with lysis buffer and four times with 50 mM NH_4_HCO_3_. Next, beads were incubated with 10 ng/µL trypsin in 1 M urea 50 mM NH_4_HCO_3_ for 30 minutes, and washed once with 50 mM NH_4_HCO_3_. Subsequently, the supernatant was digested in presence of 1mM DTT over night. Digested peptides were alkylated and desalted prior to LC-MS analysis.

### Mass Spectrometry

For the mass spectrometry analysis, the desalted peptides were separated in a 15 cm analytical column C18 micro column (75 µm ID homepacked with ReproSil-Pur C18-AQ 2.4 µm from Dr. Maisch) with a 40 min gradient from 5 to 60% acetonitrile in 0.1% formic acid (Ultimate 3000 RSLCnano system). The effluent from the HPLC was directly electro-sprayed into a LTQ-Orbitrap mass spectrometer (Thermo Fisher Scientific). The MS instrument was operated in data-dependent mode to automatically switch between full scan MS and MS/MS acquisition. Survey full scan MS spectra (from m/z 300 – 2000) were acquired in the Orbitrap with resolution R=60,000 at m/z 400 (after accumulation to a ‘target value’ of 500,000 in the linear ion trap). The six most intense peptide ions with charge states between 2 and 4 were sequentially isolated to a target value of 10,000 and fragmented in the linear ion trap by collision induced dissociation (CID). All fragmentation spectra were recorded in the LTQ part of the instrument. For all measurements with the Orbitrap detector, 3 lock-mass ions from ambient air were used for internal calibration as described before (Olsen *et al*, 2005). Typical MS conditions were: spray voltage, 1.5 kV; no sheath and auxiliary gas flow; heated capillary temperature, 200°C; normalized CID energy 35%; activation q = 0.25; activation time = 30 ms. MaxQuant 1.5.2.8 was used to identify proteins and quantify by iBAQ. Maxquant conditions were: Database, Uniprot_Spombe_150114; MS tol, 10 ppm; MS/MS tol, 0.5 Da; Peptide FDR, 0.1; Protein FDR, 0.01 Min. peptide Length, 5; Variable modifications, Oxidation (M); Fixed modifications, Carbamidomethyl (C); Peptides for protein quantitation, razor and unique; Min. peptides, 1; Min. ratio count, 2.

### Small scale protein analysis

For examining cellular protein levels by immunoblots, extracts were prepared under denaturing conditions (Knop *et al*, 1999). Co-immunoprecipitations were performed with cell extracts prepared under native conditions, similarly to the large scale protocol, either from frozen cell droplets using a freezer mill [SPEX, 6970EFM] or from frozen cell pellets using a cell homogenisator [Precellys 24, Peqlab]. Ubiquitylation pull-downs were done under denaturing conditions essentially as described (Braun *et al*, 2011), except using Ni-NTA Magnetic Agarose Beads [Qiagen, 336113]. Protein levels were analyzed by immunoblots with anti-polyHistidine [Sigma, H1029], anti-FLAG [Sigma, P3165] and anti-HA [Sigma, H6908] antibodies.

### ChIP assays

ChIP experiments were performed essentially as described (Barrales *et al*, 2016). Cross-linking was performed with 1% formaldehyde for 10 min at RT. For quantitative ChIP, immunoprecipitations were performed with 2–5 µg of antibody against H3 (or H3 modifications) or epitope-tagged proteins with lysates corresponding to 15 OD_600_ and 60 OD_600_, respectively. The following antibodies were used: anti-GFP, anti-phospho RNA pol II (Ser5) (provided by A. Ladurner, LMU), anti-Flag [Sigma, F3165], anti-HA [Sigma, H6908], anti-H3 [Active Motif, 61475], anti-H3K9me2 [Abcam, ab1220] and anti-H3K14ac [Abcam, ab52946]. Immunoprecipitated DNA was quantified by qPCR using the PowerUp(tm) SYBR Green Master mix [Life Technologies, A25778] and a 7500 Fast real-time PCR system [Applied Biosystems]. Primers are listed in Supplemental Table S3. Unless otherwise noted, mean and SEM values were calculated from three independent experiments. qPCR signals were normalized against the input samples for each primer position, which avoids any bias against subtelomeric sequences that might be under- or overrepresented due the potential shortening of telomeric ends or recombination in absence of Ccq1. For ChIP experiments with histone proteins, histone modifications, or RNA pol II, input-normalized qPCR signals were additionally normalized to the mean of 2-3 euchromatic loci (*act1*^*+*^, *ade2*^*+*^, *tef3*^*+*^) as an internal control, which was set to 1 (‘EC normalized’).

**Internal ChIP normalization**:

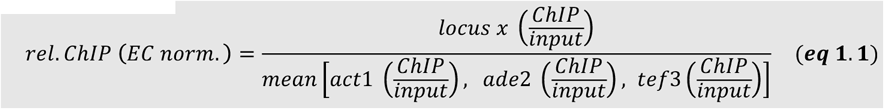

Using the mean of multiple euchromatic loci (‘EC’) instead of single locus (e.g. *act1+*) reduces bias coming from variations in ChIP experiments, especially when doing IP experiments with bulk chromatin proteins (see also Suppl. Fig S3). For ChIP-seq, anti-H3 immunoprecipitated DNA corresponding to a total of 120 OD_600_ starting material (whole cell lysate) was pooled from independent experiments. The samples were treated with 0.04 mg of RNase A (30 min, 37°C) and 0.04 mg of Proteinase K (1.5 hour, 37°C) and purified [Zymo Research, D5201]. Samples of at least three different ChIP experiments were combined and prepared for sequencing using NEBnext Ultra II DNA Library prep Kit of Illumnia [NEB, E7645S]. Single-end sequencing of libraries, demultiplexing, mapping of Illumina reads and normalization were performed as previously described (Brönner *et al*, 2017).

### RT-qPCR analyses

RT-qPCR experiments were carried out as previously described (Braun *et al*, 2011). cDNA was quantified by qPCR using primers listed in Supplemental Table S3. Prior calculation of mean and SEM values from independent experiments, *act1*^*+*^ normalized data sets were standardized to the mean of all samples from each experiment (experimental normalization; *eq 2.1*. These sample pool-normalized results were shown relative to the mean value of the sample pool-normalized wild type data from all (n) experiments (*eq 2.2*).

**experimental normalization**:

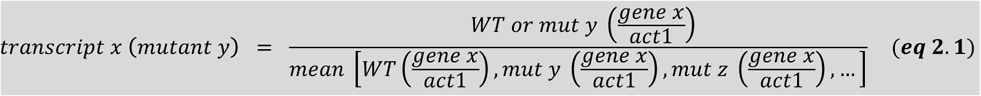

**mean WT normalization:**

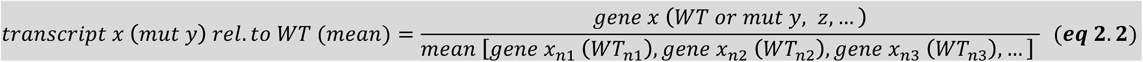

Using the average from a collection (sample pool) instead of a single strain (e.g. WT) reduces bias, especially when transcripts levels are low in the repressed state and therefore more prone to noise.

### Telomere length and genomic stability assays

Telomere-PCR was performed as previously described (Moravec *et al*, 2016). Briefly, telomeres of denatured genomic DNA were poly(C)-tailed. Telomeres were then amplified by PCR using primers that anneal to the poly(C) tail and a downstream region. Telomere length was determined by analyzing PCR products on a 0.8% agarose gel. Primers used for Telomere-PCR are shown in Table S3. For the telomere stability assay with *ccq1Δ* cells, fresh deletions mutants were generated by homologous recombination using standard genome engineering methods. Individual knock-out clones grown for 5 days on selective media were used to inoculate 5 mL of media at OD_600_ ∼ 0.05. Cultures were grown over night to OD_600_ = 3-5. Amount of cells corresponding to 5 OD_600_ was harvested by centrifugation and flash-frozen. Remaining culture was back diluted to OD_600_ ∼ 0.05 to be grown 24 hours. This process was repeated to collect samples from 6 days. Genomic DNA was extracted from frozen cell pellets using a yeast DNA extraction kit [Thermo Fisher Scientific, 78870]. Relative copy numbers of individual reporter genes or subtelomeric loci were measured by qPCR and normalized to internal reference genes (*act1*^*+*^, *ade2*^*+*^, *tef3*^*+*^) as described above. For quantifying the minichromosome loss of *Ch16 m23::ura4* (Suppl. Fig. S8), the genomic copy number of the *ade6*^*+*^ gene was quantified by qPCR using oligonucleotides that anneal to both the chromosomal *ade6-M210* allele (Chr III) and the *ade6-M216* allele (Ch16), followed by normalization to reference genes (EC: *act1*^*+*^, *ade2*^*+*^, *tef3*^*+*^). Based on the rationale that the level of Ch16 and *m23::ura4*^*+*^ (encoded by the minichromosome) are equal at the start of the experiment, the relative level of *ade6*-*M216*_*Ch16*_ allele was determined by subtracting the copy number of *ura4*^*+*^ at day 1 from the total copy number of *ade6*^*+*^ (i.e. *ade6*-*M216*_*Ch16*_ + *ade6-M210*_*ChrIII*_) for each time point. These calculated *ade6-M216*_*Ch16*_ values were used for normalizing the *m23::ura4*^*+*^ copy number.

## ACKNOWLEDGEMENTS

We thank Bassem Al-Sady, Ramón Ramos Barrales and members of our lab for critical reading of the manuscript, and Guy Riddihough (Life Science Editors) for editorial assistance. We thank Robin Allshire, Zac Cande, Songtao Jia, Junko Kanoh, Andreas Ladurner, Michaela Smolle, Philipp Korber, and Takashi Toda for strains, plasmids and reagents. We also thank Sebastian Eustermann and the Hopfner Lab (Gene center, LMU) for technical training and sharing equipment, and Scott Wood for initial efforts to identify CLRC interactors. We further thank Songtao Jia for communicating results prior to publication. This work was supported by grants awarded to S.B. from the German Research Foundation (BR 3511/2-1) and the European Union Network of Excellence EpiGeneSys (HEALTH-2010-257082). S.B. is Member of the Collaborative Research Center 1064 funded by the German Research Foundation and acknowledges infrastructure support.

## AUTHOR CONTRIBUTIONS

TvE, MF, and SB designed the study. MF, TvE, MC and SFB generated plasmids and yeast strains. MF performed biochemical purifications, pull-down experiments and immunoblots. IF designed and performed the LC-MS analysis under supervision by AI. MF and SFB carried out yeast growth assays. The UBA trap method was developed by MS in the lab of DT. TvE, MF, SFB, ZS, MC, LMC and SB performed RT-qPCR and ChIP experiments. TvE conducted genome stability and telo-PCR assays. TvE performed ChIP-seq together with CB and MH. TvE, MF, IF, CB, ZS, MC, LMC and SB carried out data analysis. SB, TvE, and MF wrote the manuscript and IF and AI contributed to editing.

---

**Figure S1:**
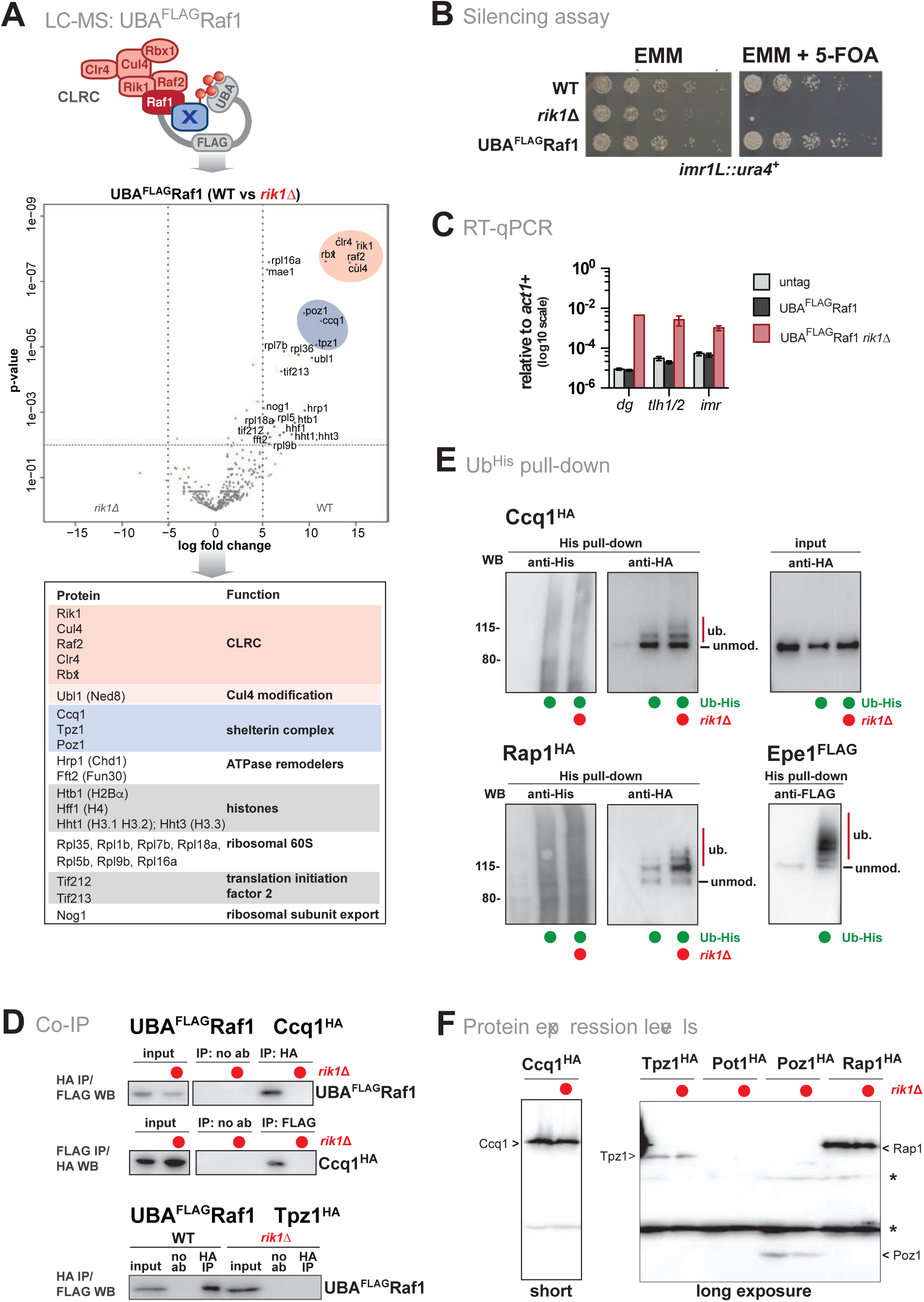
(Supplement to Figure 1). CLRC interacts with shelterin without affecting ubiquitylation. **(A)** Mass spectrometry analysis of proteins co-purified with Raf1 fused to the ubiquitin-associated (UBA) domain of the ubiquitin receptor Dsk2 from *S. cerevisiae* (UBA^FLAG^Raf1). This method is based on the *Ubiquitin Ligase Trapping* method originally developed in *S. cerevisiae* (Mark *et al*, 2013), which stabilizes the binding of substrates to the ubiquitin ligase. The volcano plot depicts proteins enriched in UBA^FLAG^Raf1 WT cells relative to UBA^FLAG^Raf1 *rik1*Δ. Members of the CLRC and shelterin complexes are highlighted in red and blue, respectively. Bottom panel displays nuclear proteins enriched (log_2_ ≥ 5 or pval ≤ 0.01). **(B)** Silencing reporter assay at *imr1L::ura4*^*+*^. Fivefold serial dilution on non-selective (EMM) and selective media (EMM+5-FOA) of WT, *rikΔ* and UBA^FLAG^Raf1 cells. The N-terminal fusion does not interfere with the function of CLRC in heterochromatin formation. The *rik1Δ* strain was used as negative control. **(C)** RT-qPCR analysis in strains expressing untagged Raf1, UBA^FLAG^Raf1 and UBA^FLAG^Raf1 *rik1Δ*. Values are shown as mean values ± SEM from 3 independent experiments. **(D)** Co-immunoprecipitation of Raf1 with Ccq1 (top and middle panel) and Tpz1 (bottom panel) in presence and absence of Rik1. Strains expressing endogenous levels of UBA^FLAG^Raf1 and Ccq1^HA^ or Tpz1^HA^ were subjected to anti-HA immunoprecipitation in the presence of Benzonase (indicating that the interactions are independent of DNA or RNA; negative control: no antibody, noAb). Input and immunoprecipitated material were analyzed by anti-FLAG immunoblots. For Ccq1^HA^ the reciprocal experiment is also shown. **(E)** Ubiquitin pull-down experiments. Top panels shows *in vivo* ubiquitylation of Ccq1^HA^ in WT and *rik1Δ* cells expressing 6His-ubiquitin. Left panels: precipitated 6His-ubiquitin conjugates (20%) analyzed by anti-His and anti-HA immunoblots. Right panel: input fraction (0.1%) analyzed by anti-HA immunoblot. The bottom panels show 6His-ubiquitin pull-down assays for Rap1^HA^ in WT and *rik1Δ* cells (left) and Epe1^FLAG^ in WT cells (right). Epe1 is shown as an example for poly-ubiquitylation. WT cells expressing untagged ubiquitin are used as negative control. **(F)** Anti-HA immunoblot of epitope-tagged shelterin proteins expressed from their endogenous locus in WT and *rik1Δ* cells. The asterisks denote unspecific signals derived from cross-reactions with the anti-HA antibody. Note that Ccq1 is expressed at higher levels compared to other shelterin subunits; endogenous levels of Pot1 are low and not detected by the immunoblot. Estimated sizes (kDa) of the tagged proteins: Ccq1 (86 kDa), Tpz1 (61 kDa), Pot1 (67 kDa), Poz1 (34 kDa) and Rap1 (84 kDa).

**Figure S2:**
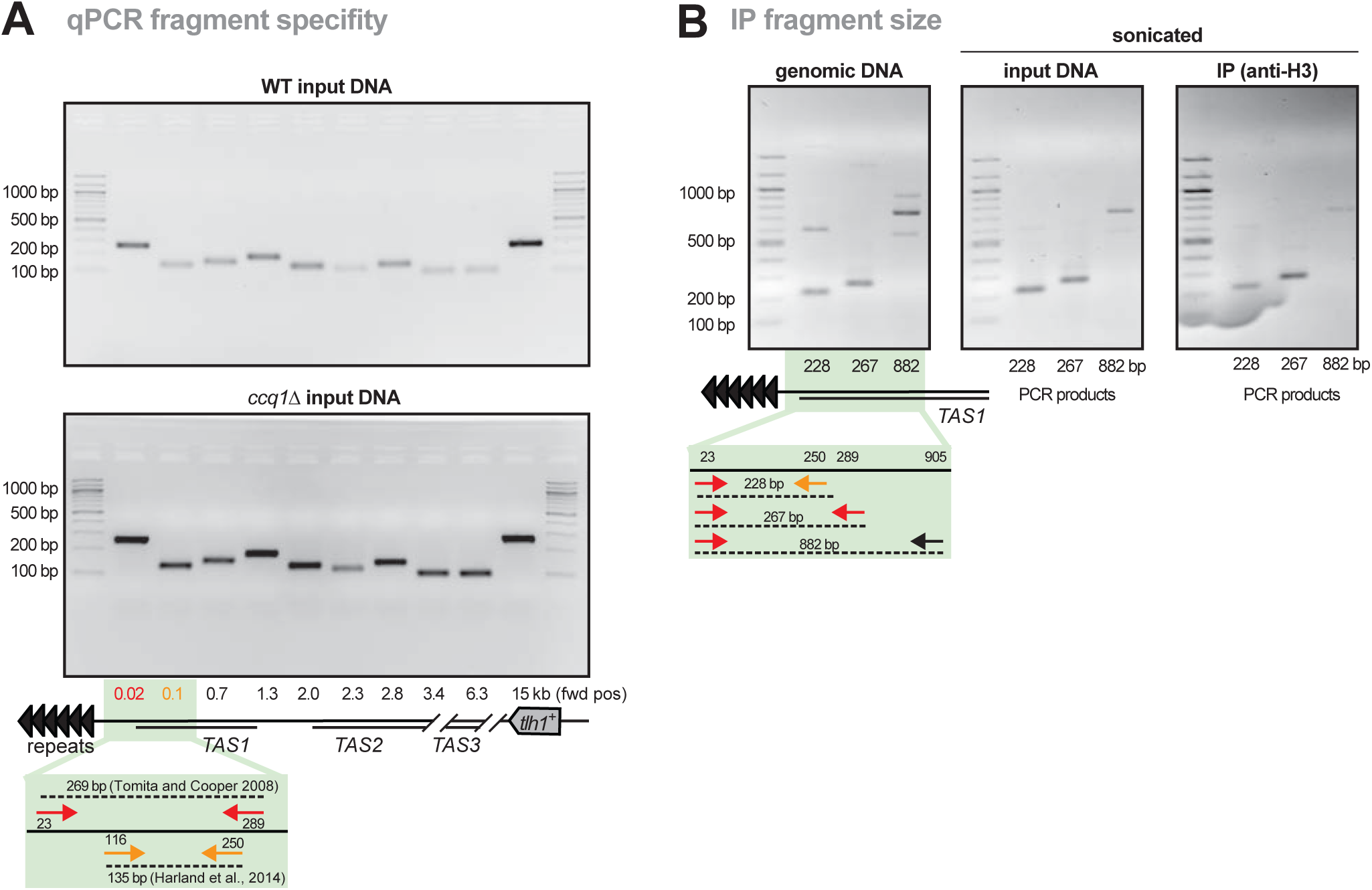
(Supplement to Figure 1). qPCR primer specificity and DNA fragment size from ChIP samples. **(A)** qPCR products of indicated primer pairs using WT and *ccq1Δ* genomic DNA (input material from ChIP reactions) were analyzed on a 2% agarose gel. Each primer pair results in a single product and reproducible product size for both WT and *ccq1Δ* samples, ruling out the possibility that potential genetic rearrangements upon deletion of *ccq1*^*+*^ affect primer specificity. The most telomere-proximal primers were used in previous studies (Tomita & Cooper, 2008; Harland et al, 2014). Note that while the ‘forward’ primers of these primer pairs anneal to different regions (i.e. 23 bp and 116 bp relative to the telomeric repeats, here denotated as 0.02 kb and 0.1 kb), the resulting PCR products overlap due to the position of the corresponding ‘reverse’ primers (i.e. 289 bp and 250 bp, respectively). Nonetheless, ChIP experiments performed with Shelterin, CLRC and H3K9me2 (see Figure 1C-E, S3A-B, 2B, respectively) suggest that these primer pairs appear to discriminate between different regions. **(B)** DNA fragment sizes of telomere-proximal fragments (TAS1) DNA analyzed with different primer combinations, as indicated in the scheme. Shown are results for genomic DNA (not-sonicated, left), input DNA (sonicated, middle) and H3 ChIP DNA (sonicated, right). Note that despite multiple primer binding sites (due to the repetitive nature of the TAS) sonication suppresses the appearance of additional (unspecific) PCR products and that products longer than 800 bp are hardly detected in the ChIP material.

**Figure S3.**
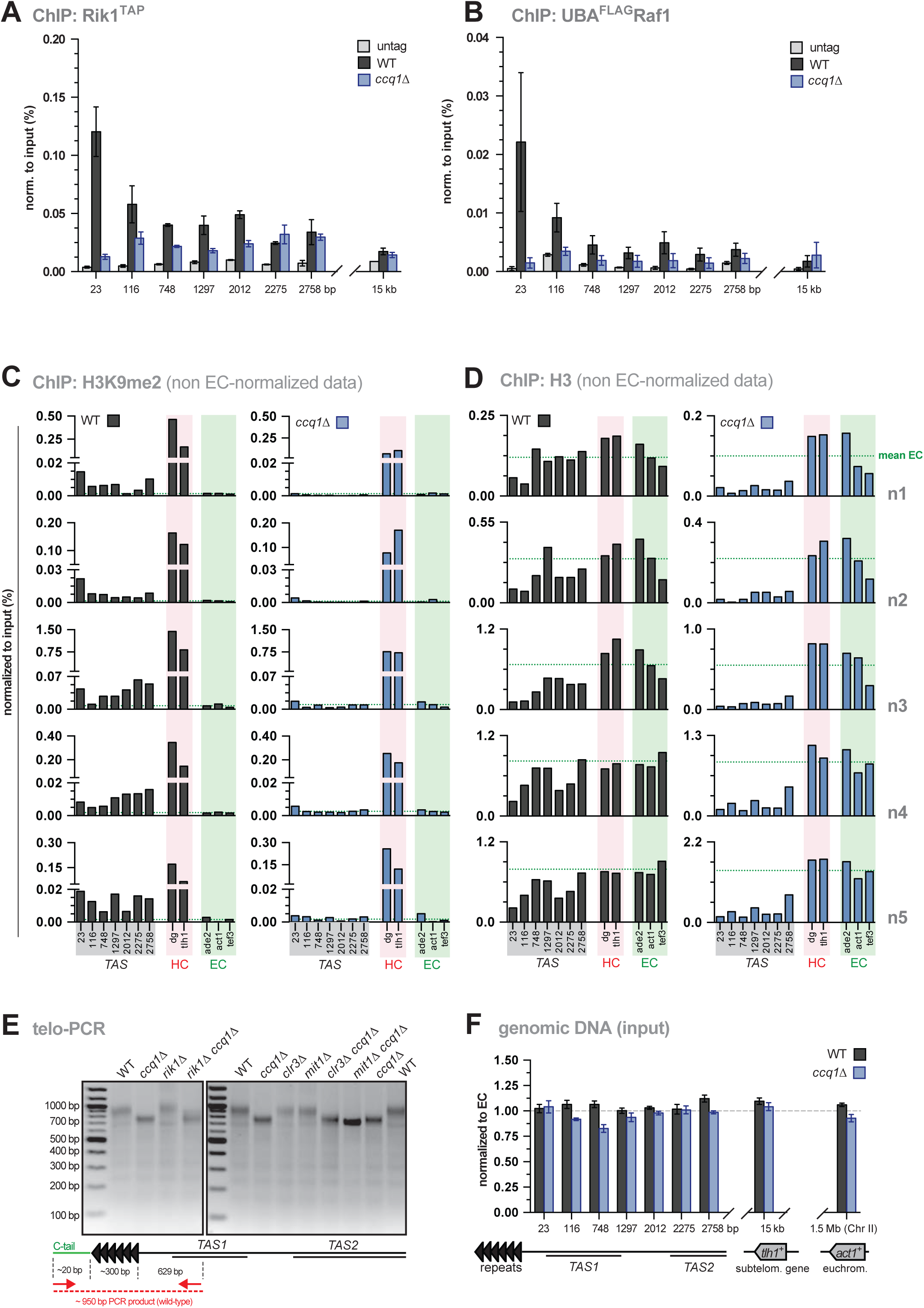
(Supplement to Figure 2). Normalization of histone qPCR-ChIP data and telomere length detection in WT and (early) *ccq1Δ* cells. **(A,B)** ChIP-qPCR analysis of Rik1^TAP^ (A) and UBA^FLAG^Raf1 levels (B) in WT and *ccq1*Δ cells (negative control: untagged strain). ChIP data are normalized to input and shown are mean values from 2 and 3 independent biological experiments (error bars: range and SEM) for Rik1^TAP^ and UBA^FLAG^Raf1, respectively. **(C,D)** ChIP-qPCR analyses of H3K9me2 (A) and H3 (B) in WT and *ccq1*Δ cells for TAS regions, heterochromatic (HC, shaded in red) and euchromatic loci (EC, shaded in green). The dotted line indicates average EC level based on levels of three euchromatic genes (*ade2*^*+*^, *act1*^*+*^ *and tef3*^*+*^). Shown are 5 independent biological replicates for WT and *ccq1Δ*. Note that relative differences among various loci are highly reproducible within an individual replicate despite some variability in ChIP efficiency among different replicates, underscoring the significance of using internal references for ChIP normalization. **(E)** Telomere length analysis of different mutants using telo-PCR, which amplifies the entire repeats and 629 bp of the adjacent TAS1 region (Moravec *et al*, 2016). PCR products are analyzed on a 0.8% agarose gel. Note that telomeres are shortened in early *ccq1Δ* (∼200 nt) but still retain telomeric repeats. Telomeric shortening is not seen in mutants deficient in CLRC or SHREC despite the similar nucleosome-loss phenotype as seen in *ccq1Δ* cells (see Fig. 3). **(F)** qPCR analysis of genomic input DNA in WT and *ccq1*Δ cells normalized to EC which is set to 1 (dotted line). Only minor differences (i.e. < 20%) were detected between WT and *ccq1Δ*, which have been taken into account by normalizing ChIP data (input normalization; see methods). qPCR data are represented as mean ± SEM from 3 independent experiments.

**Figure S4:**
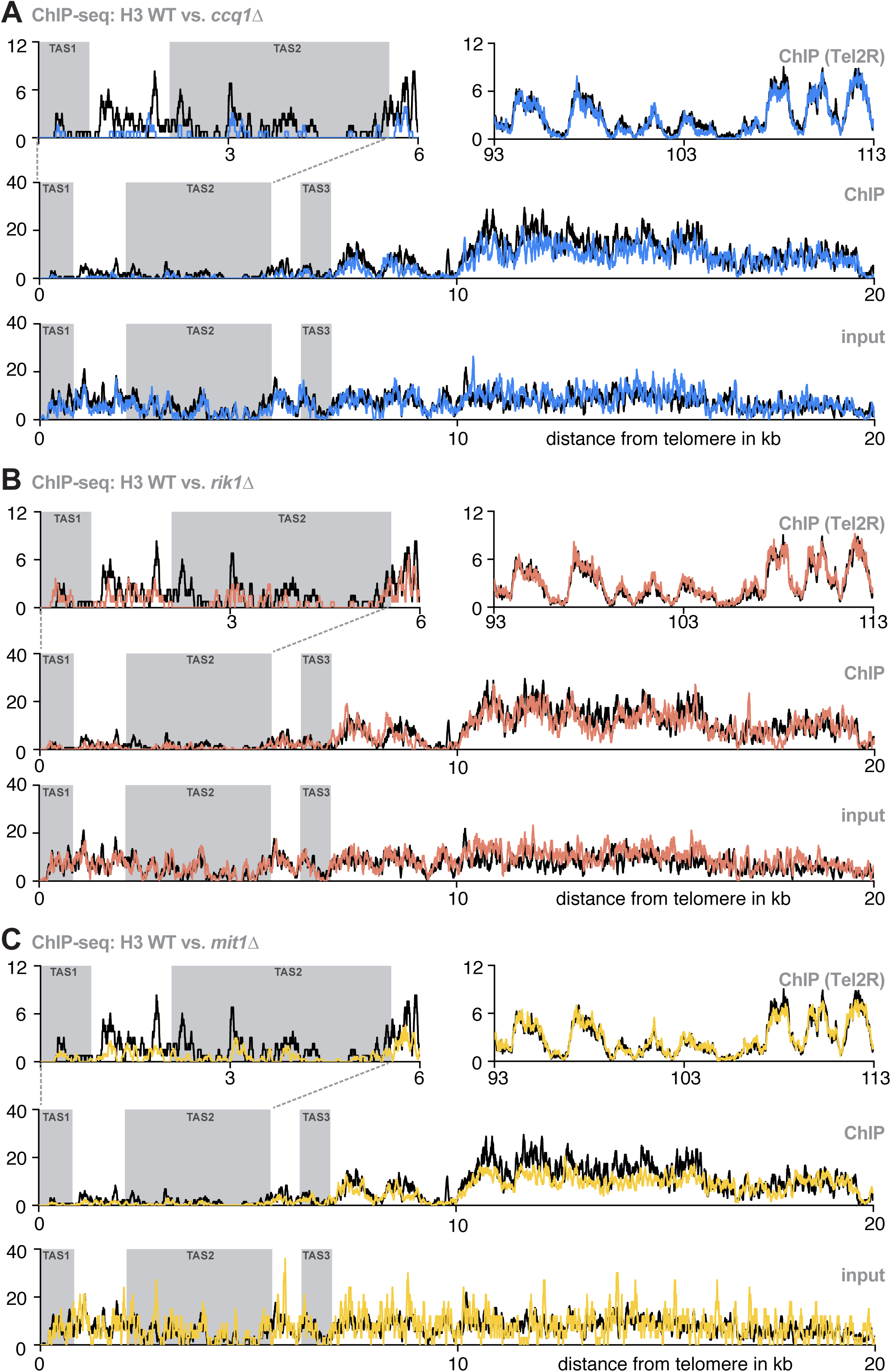
(Supplement to Figure 4). Subtelomeric H3 profiles in WT cells and mutants deficient in Ccq1, CLRC, and SHREC. **(A-C)** ChIP-seq reads mapped to TEL2R (shown in reverse orientation for consistency) for WT (black) vs. *ccq1*Δ (blue) (A), *rik1*Δ (red) (B) and *mit1*Δ (yellow) cells (C). Shown are reads of ChIP data for TAS1 and TAS2 (top left panel), a nucleosome-free region reported by Tashiro et al., 2017 (Tashiro *et al*, 2017) (top right panel), and the subtelomeres including the TAS1-3 and *tlh1*^*+*^ (middle panel). The bottom panel shows input samples for the same region. Grey shaded areas are TAS regions.

**Figure S5.**
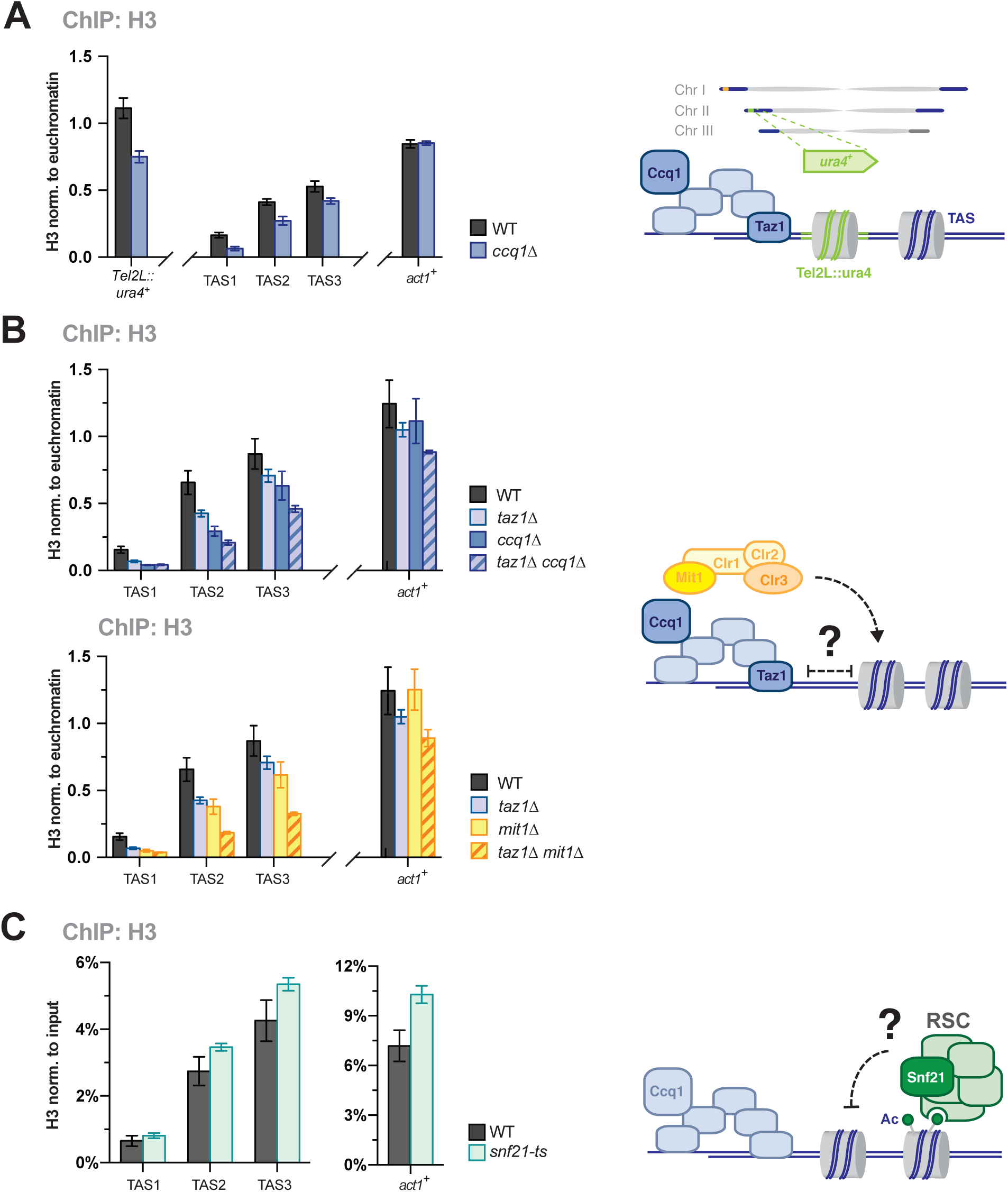
(Supplement to Figure 5). Low nucleosome occupancy at TAS is not caused by proximity of telomeric repeats or shelterin binding, RSC. **(A)** ChIP-qPCR analysis of H3 at reporter gene (*ura4*^*+*^) and TAS regions WT and *ccq1Δ* strains (see scheme). **(B)** ChIP-qPCR analysis of H3 in WT, *ccq1Δ*, and *taz1Δ* (top) or *mit1Δ* (bottom) and corresponding double mutants (see scheme). For A and B, data are represented as mean ± SEM from 3 independent experiments (normalization as in Fig. 2B). **(C)** ChIP-qPCR analysis of H3 in WT and *snf21-ts* cells (see scheme). Data are normalized to input and represented as mean ± SEM from 3 independent experiments (each derived from 2-3 parallel ChIP samples). Note that the global role of RSC precludes using internal controls (i.e. EC) for normalization.

**Figure S6:**
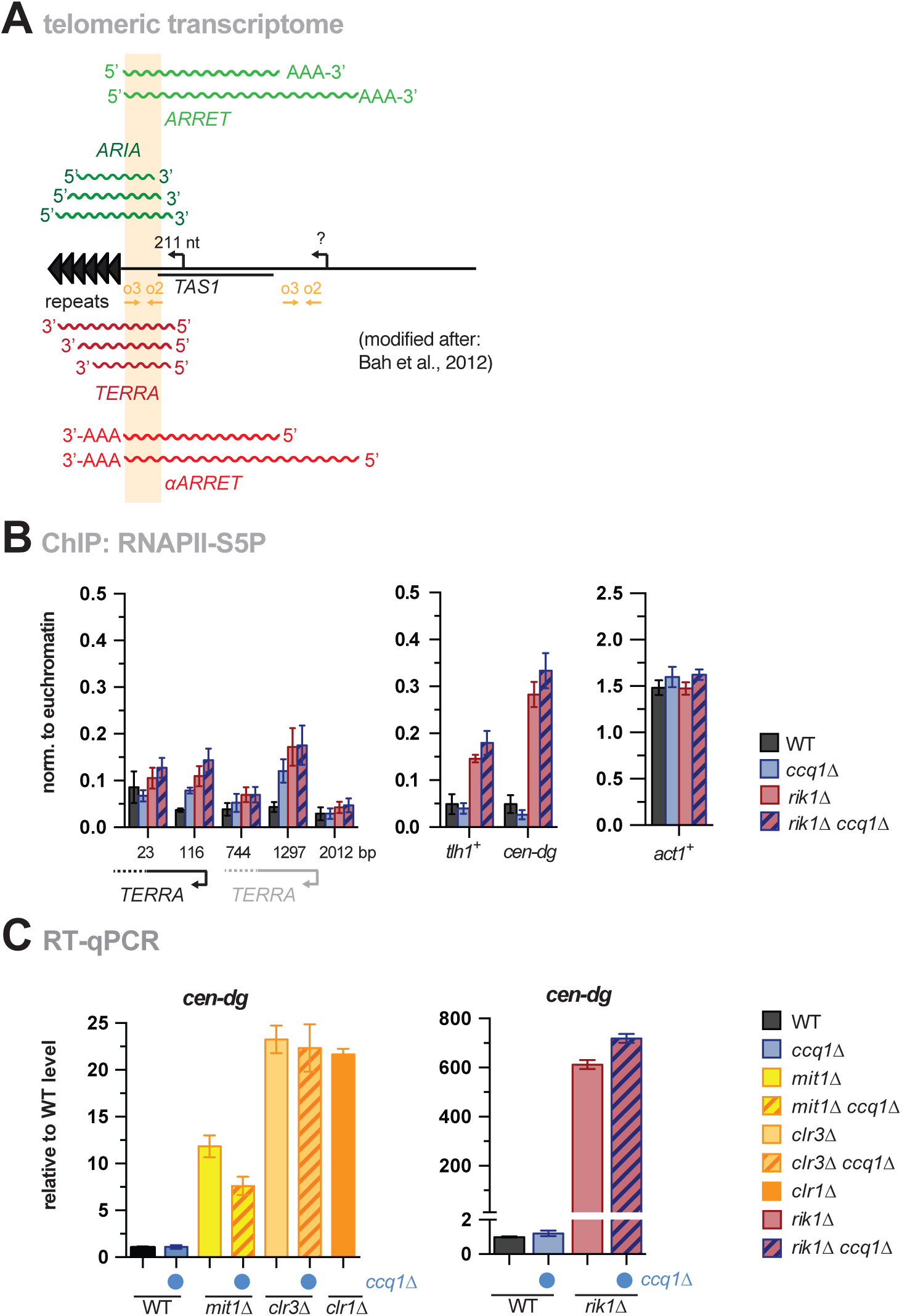
(Supplement to Figure 6). Expression of telomeric transcripts in the absence of CLRC and SHREC. **(A)** Scheme of telomeric transcripts (modified after (Bah *et al,* 2012)). Whereas *ARRET* and *αARRET* RNAs have a poly-A tail, only a small percentage of *TERRA* and *ARIA* transcripts are poly-adenylated. The amplicon (primers o2 + o3) used for RT- and ChIP-qPCR analysis anneals to all telomeric transcripts without discriminating strand specificity or shorter species that lack transcribed parts from the telomeric repeats. For simplicity, we refer to these transcripts as *‘TERRA’*. An identical sequence of the o2/o3 amplicon is present in a telomere-distal region but it is unknown whether this region also contains transcription start sites. **(B)**ChIP-qPCR analysis of RNAPII-S5P in WT, *ccq1Δ, rik1Δ* and the corresponding double mutant. Data are shown relative to the average of three euchromatic genes (EC) and represented as mean ± SEM from 3 independent experiments. **(C)** Ccq1 is not involved in silencing of pericentromeric heterochromatin. RT-qPCR analysis of transcript levels of *cen-dg* in WT, *ccq1Δ, mit1Δ, clr3Δ, rik1Δ*, double mutants with *ccq1Δ* (indicated by blue dot) and *clr1Δ* cells. Data are represented as mean ± SEM from 3 independent experiments and shown relative to WT level.

**Figure S7.**
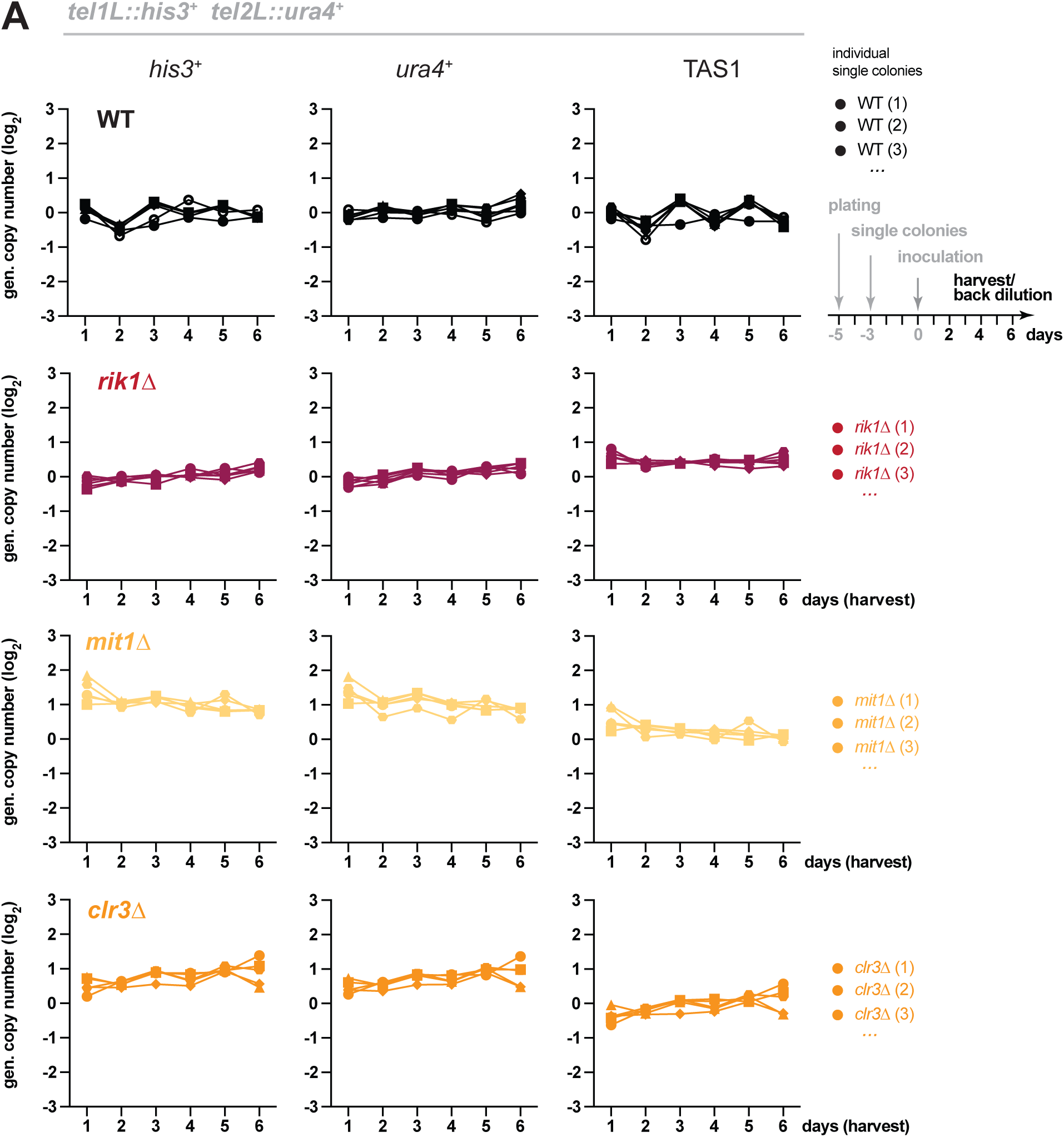
(Supplement to Figure 7). Subtelomeres do not undergo recombination in the absence of CLRC and SHREC. **(A)** Genomic copy number of *his3*^*+*^, *ura4*^*+*^ and TAS1 in indicated strains harboring the reporter genes *tel1L::his3*^*+*^ and *tel2L::ura4*^*+*^ cells. Cultures from individual WT and knockout clones (WT, n = 6; *rik1Δ*, n = 6; *mit1Δ*, n = 5; *clr3Δ*, n= 5) were inoculated at day 0 to grow in liquid media with regular back-dilution (every 24 hours, approximately 7 generations). Samples were taken at indicated harvest times, and relative copy numbers of genomic regions were assessed by qPCR (normalization against intrachromosomal loci).

**Figure S8:**
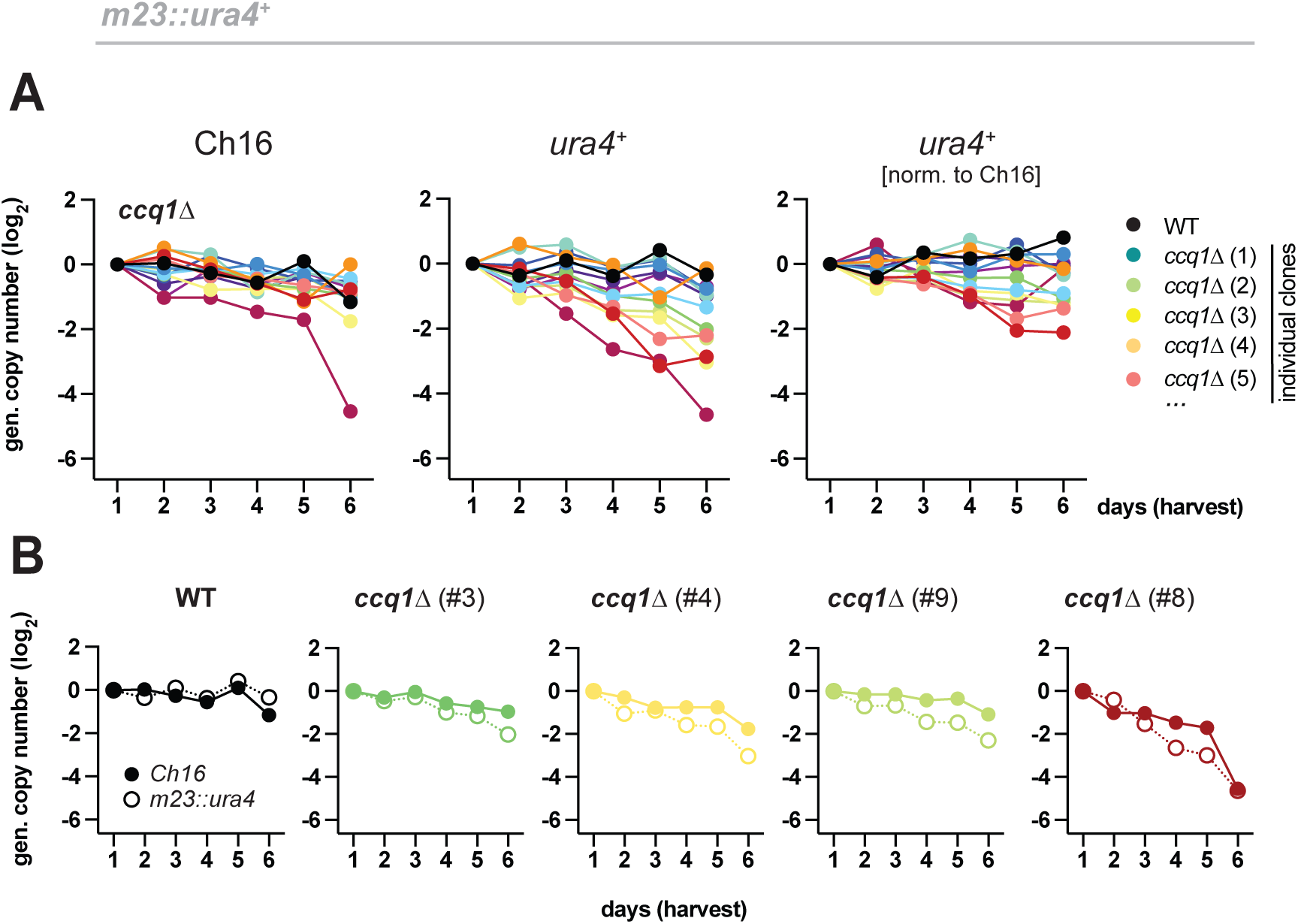
(Supplement to Figure 7). Normalization for minichromosome loss. **(A)** qPCR analysis as in (Fig. 7A) but with strains harboring the minichromosomesome Ch16 *m23::ura4*^*+*^. Right panel shows same data as Fig. 7B (note different scale); left and middle panels show analyses for minichromosome (Ch16) and *ura4*^*+*^(not normalized for minichromosome loss). (B) Selected mutants from A, displayed individually (matching color code) for direct comparison of Ch16 (closed dots) and *ura4*^*+*^ (open dots) levels.

**Supplementary table S1:**
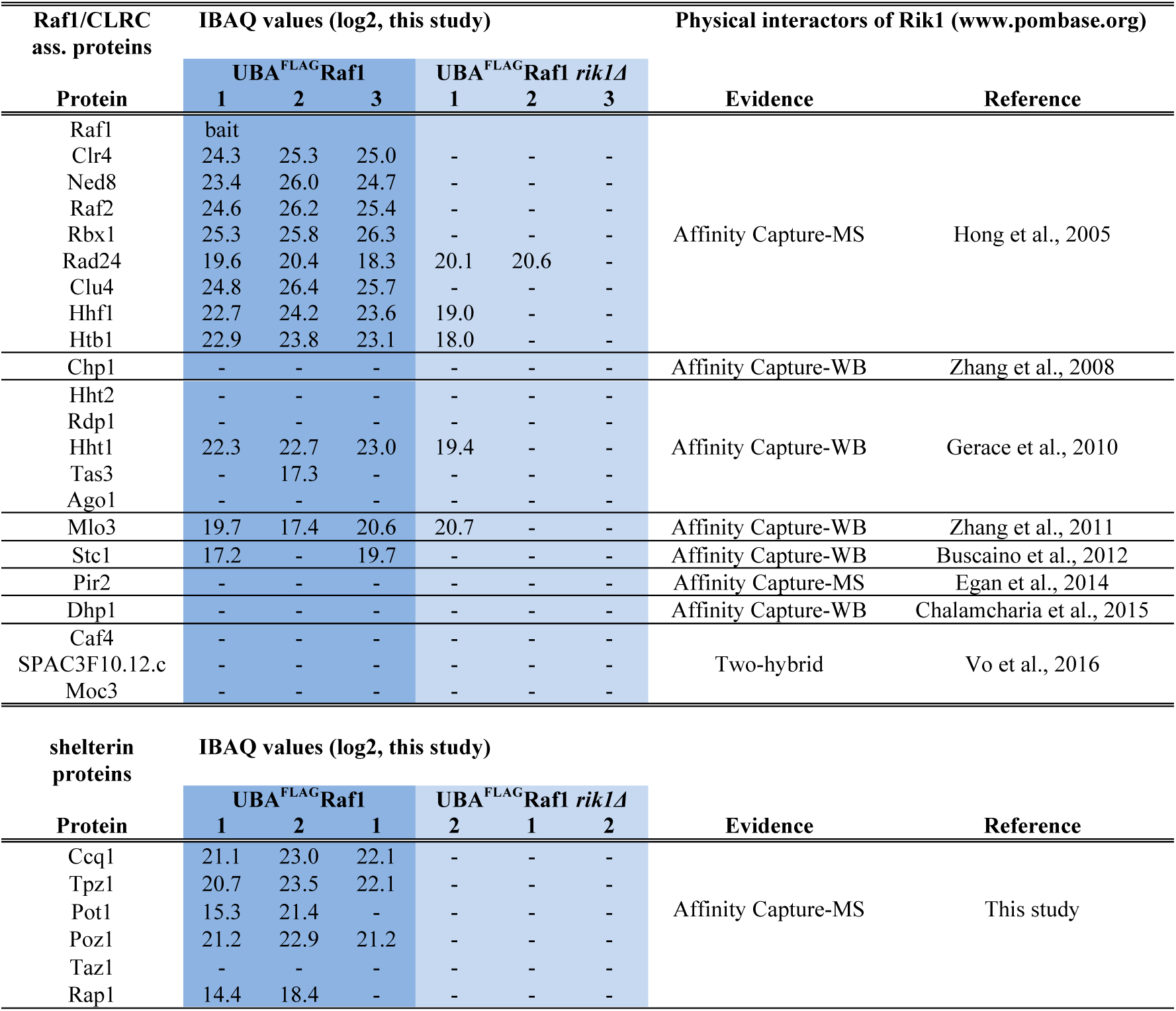
UBA^FLAG^Raf1 interactome

**Supplementary table S2:**
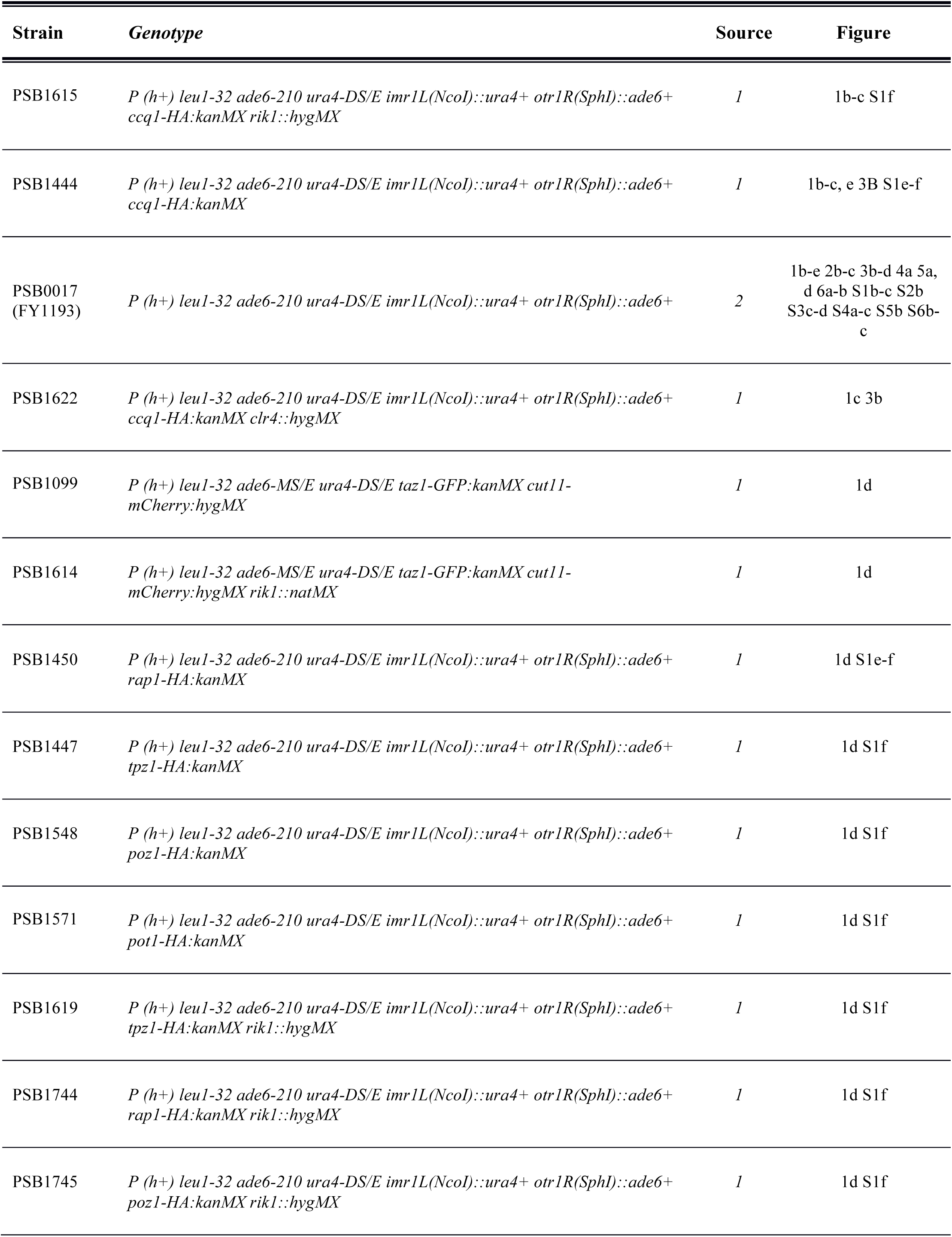
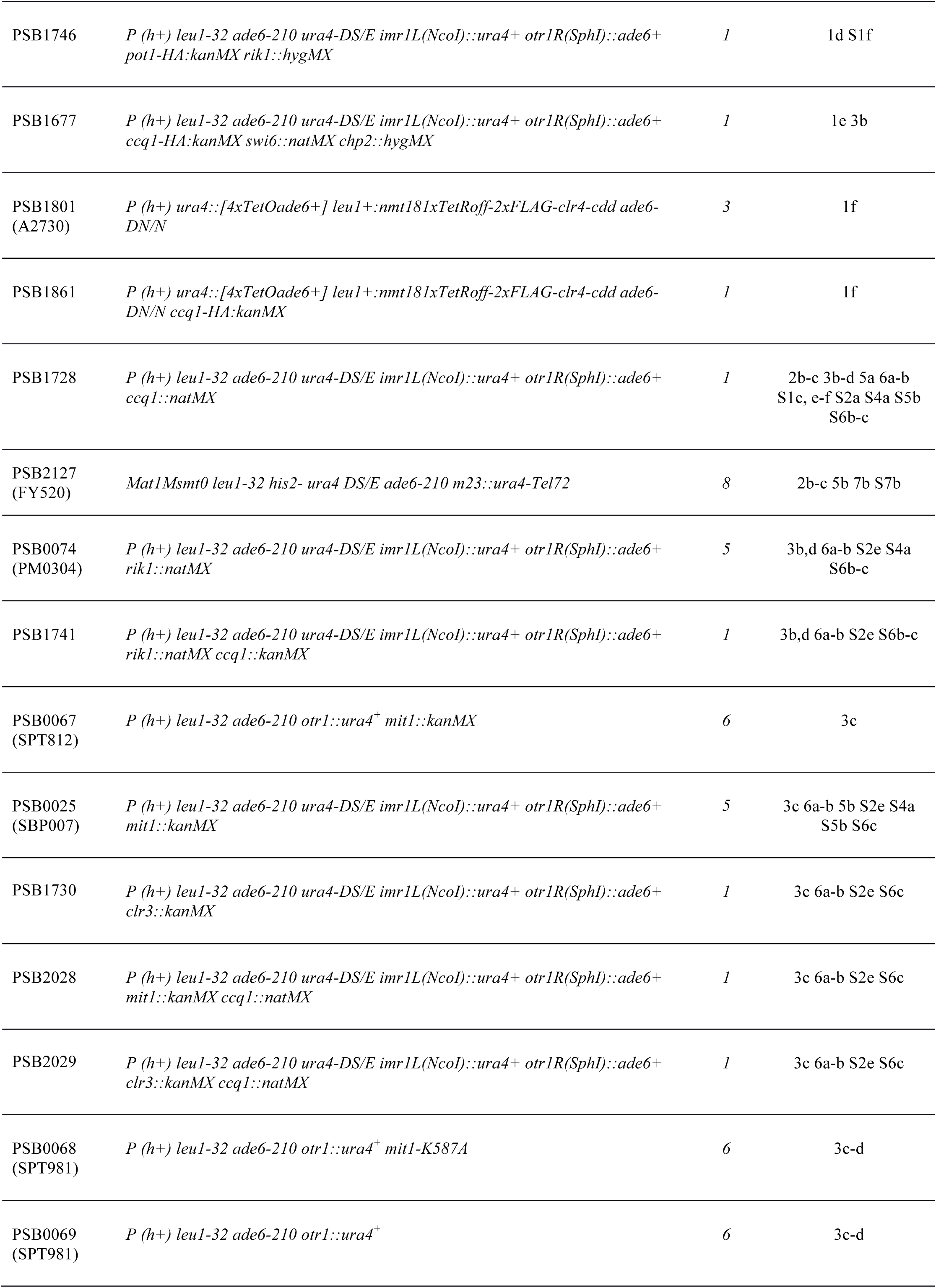
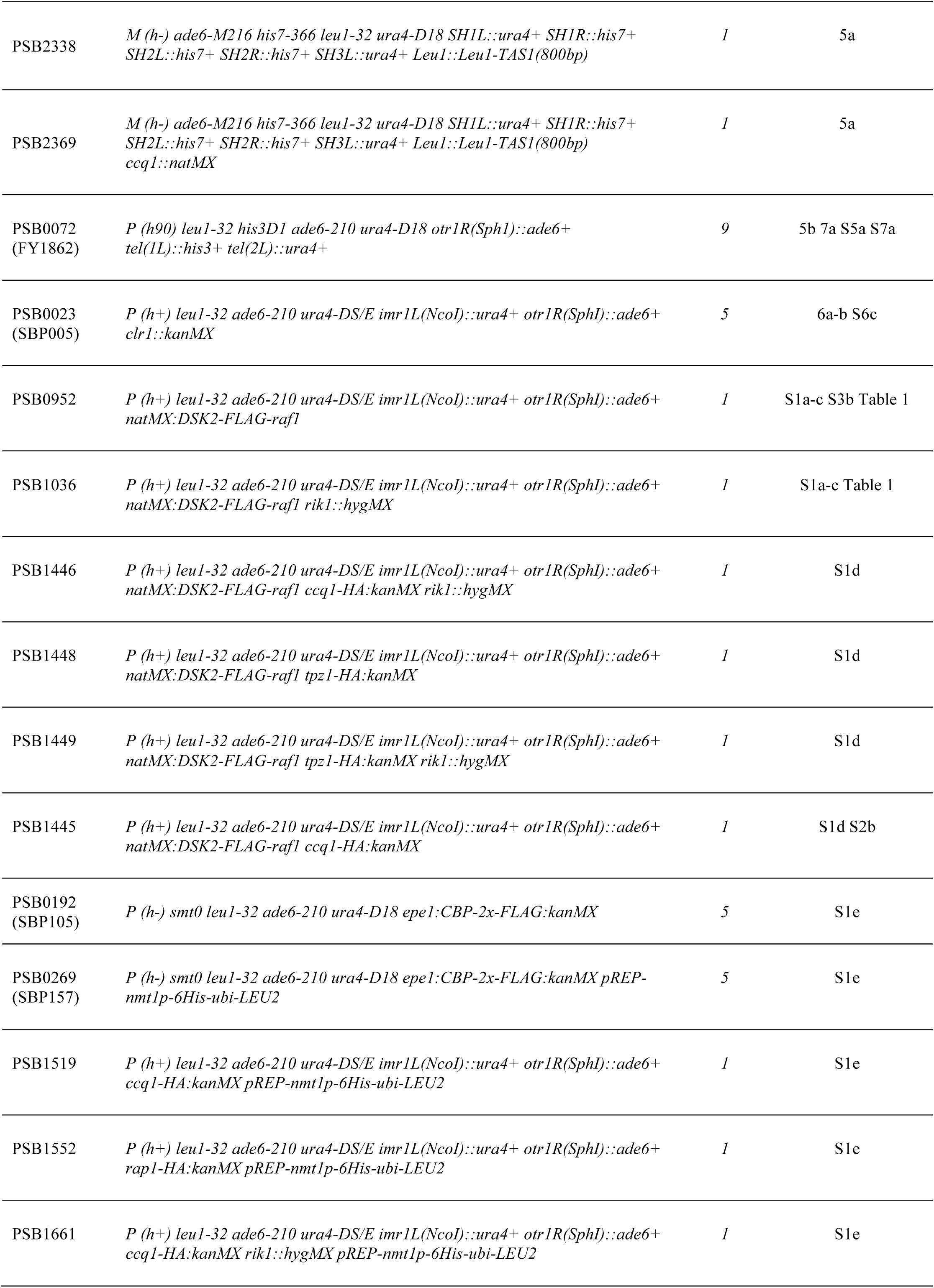
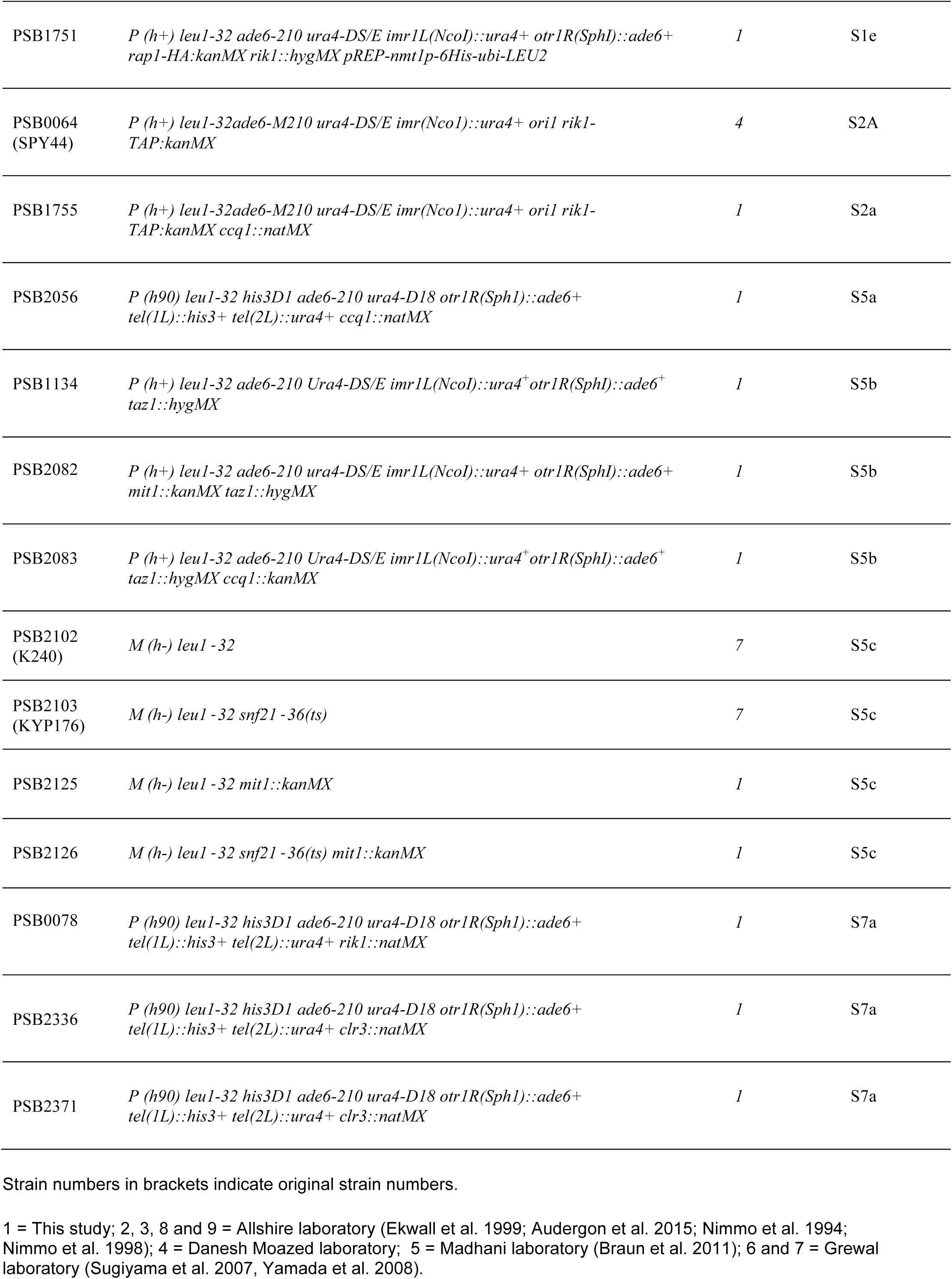
Strains used in this study, Related to Experimental Procedures

**Supplementary table S3:**
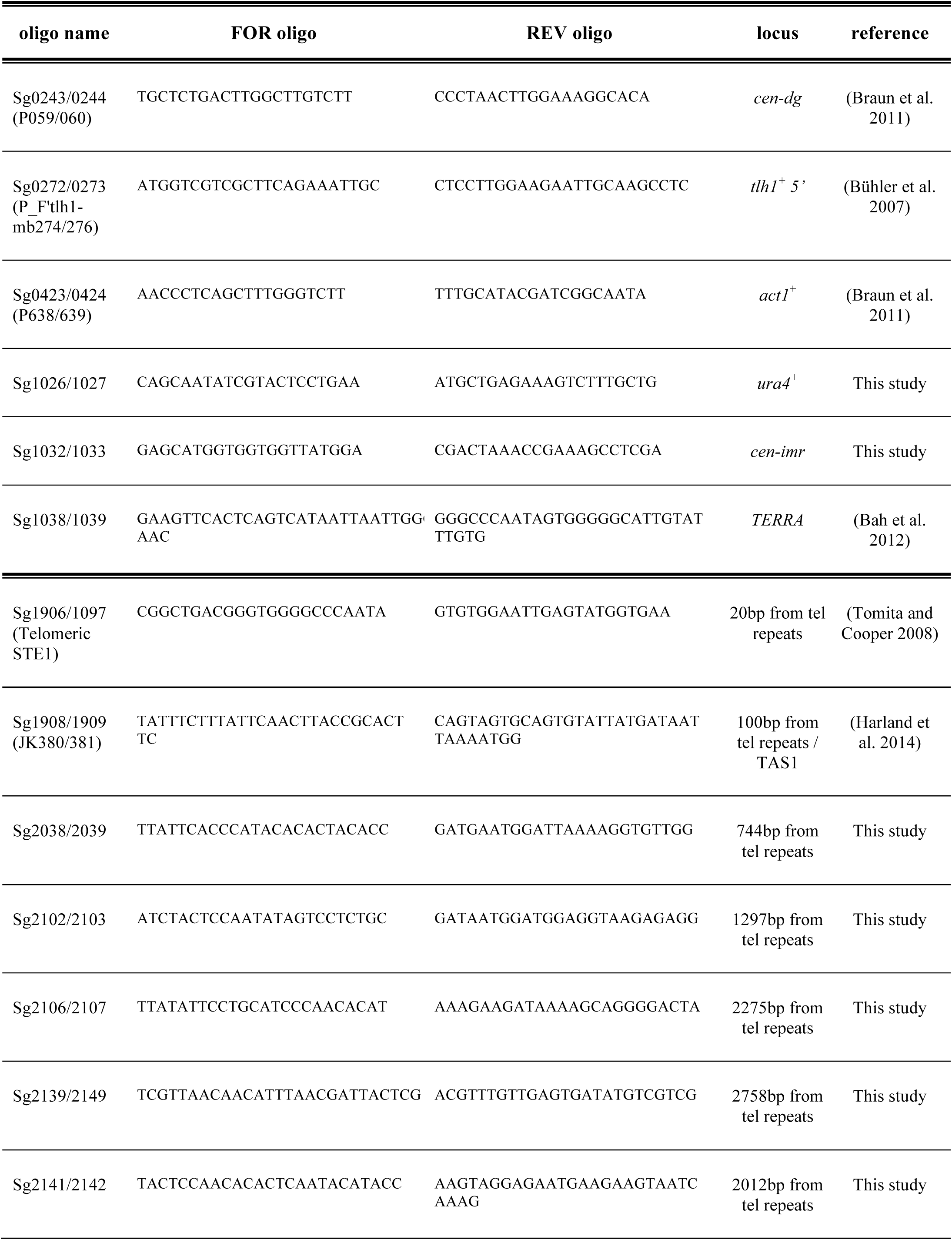
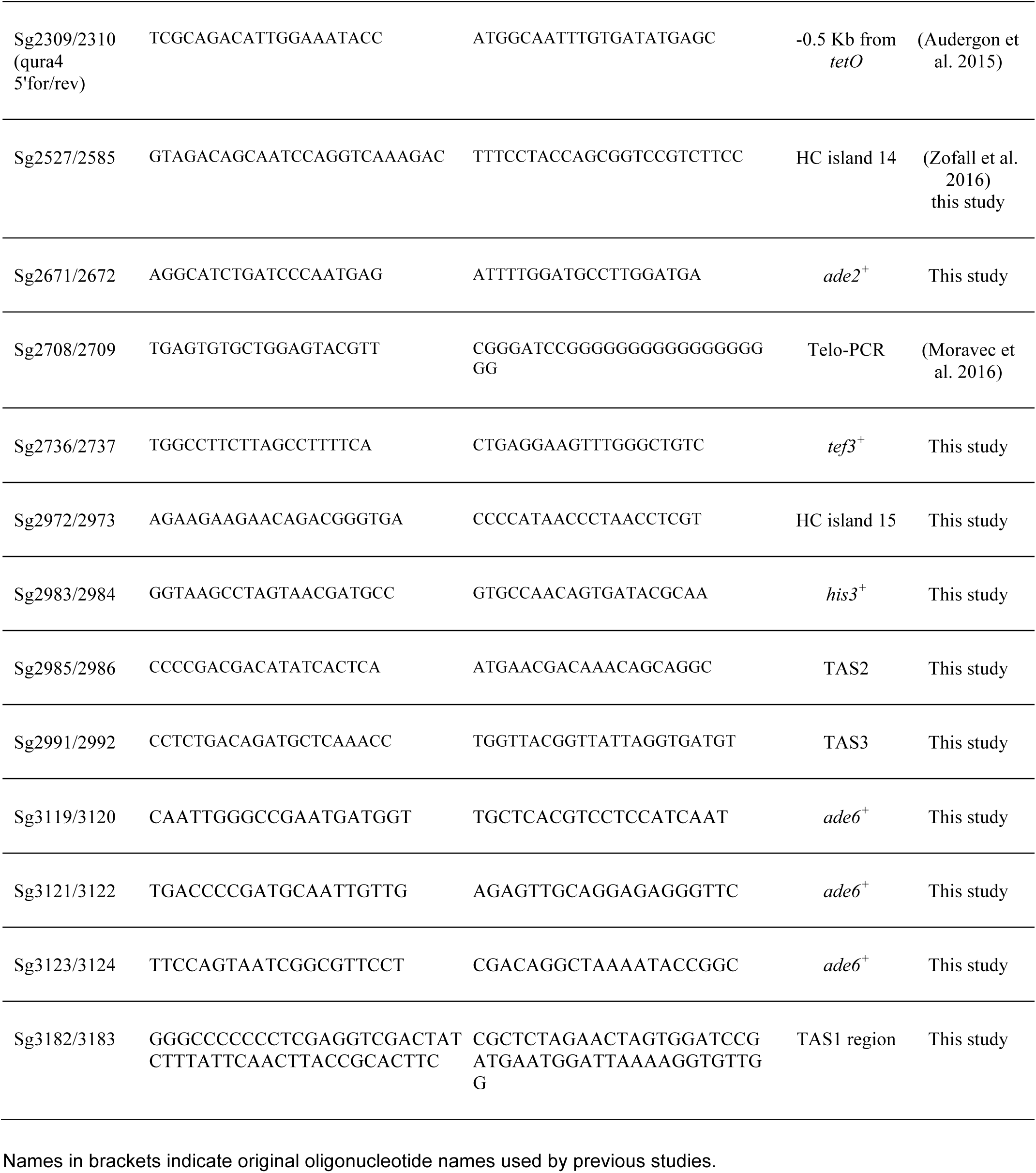
Primer sets used for RT-qPCR, ChIP analysis, Telomere-PCR and strain generation, Related to Experimental Procedures

